# Transdermal Electrophysiological Recordings of Diabetic Small Fiber Peripheral Neuropathy Using a Needle Electrode Array in Mice and Men

**DOI:** 10.1101/2023.03.03.530993

**Authors:** Magdalena Blaszkiewicz, Lydia Caron, Brooke Villinski, Joshua Passarelli, Julia M. Towne, Naeemah M. Story, Erin Merchant, Furrukh S. Khan, Nuri Emanetoglu, Leonard Kass, Rosemary L. Smith, Kristy L. Townsend

## Abstract

Diabetic peripheral neuropathy (DPN) is a common complication of diabetes. Proactive treatment options remain limited, which is exacerbated by a lack of sensitive and convenient diagnostics, especially early in disease progression or specifically to assess small fiber neuropathy (SFN), the loss of distal small diameter axons that innervate tissues and organs. We designed, fabricated, tested, and validated a first-of-its-kind medical diagnostic device for the functional assessment of transdermal small fiber nerve activity. This device, the Detecting Early Neuropathy (DEN), is an electrically conductive needle array designed to record nerve electrical activity in the skin and subdermal tissues. DEN recordings were validated across a time course of diet-induced DPN in mice, using statistical and computational analyses compared to other SFN measures. Based on these preclinical mouse data, the device design was adapted to obtain recordings in human with a flexible printed circuit board to mold to the leg or other skin regions. The DEN successfully recorded various types of neural activity in mouse and human, with or without stimulation, including validated action potentials and electromyography signals. New functional diagnostic tools like DEN offer a promising outlook for patients needing an earlier or more sensitive diagnosis of SFN/DPN, to allow for earlier and more effective treatment options, especially as more become available in the clinic in future years.

## Introduction

An estimated 30 million Americans suffer from a progressive and debilitating disease known as peripheral neuropathy (PN), the most common neurodegenerative disease^1^. More than 30 medical conditions can lead to this disease, which is often marked by a progressive loss of distal nerve endings that innervate tissues and organs. With diabetic peripheral neuropathy (DPN), the number one cause of PN, the disease typically starts in the skin of the extremities (arms and legs), and moves proximally and inward to deeper tissue layers over time. Many cases of PN are a polyneuropathy with several nerve subtypes affected (sensory, autonomic, motor), which subsequently leads to complex symptoms ranging from pain, tingling, and burning, to loss of sensation, numbness, loss of motor control, and in severe cases even limb amputation^2^. These drastic clinical outcomes are often due to a lack of effective diagnostic tools to provide an early and sensitive diagnosis of neuropathy, which could impact the clinical course of care. Currently, there is no cure for peripheral neuropathy to halt nerve degeneration and re-innervate tissues through axonal regeneration, and thus treatments for PN now primarily focus on management of symptoms (mainly analgesics) and palliative care. However, for certain peripheral neuropathies a timely diagnosis can enable interventions to prevent further decline^3^, such for DPN, where better glucose control or management of metabolic disease (such as weight loss) can be achieved to mitigate the progression of PN. DPN is a form of small fiber neuropathy (SFN), impacting the small diameter axons innervating tissues and organs. SFNs also include those caused by aging, chemotherapy, and Long COVID – all the top causes of PN overall. Currently, most functional measures of PN specifically target *large fiber nerves*, and therefore are not accurate for diagnosing SFN. Since peripheral nerves can uniquely re-grow compared to nerves in the brain (such as after nerve injuries), better functional diagnostics may provide earlier windows for interventions and opportunities for new treatments that target tissue re-innervation. This is especially important now, as more options for PN treatments are now entering clinical trials.

### Diabetic Peripheral Neuropathy (DPN) and Small Fiber Neuropathy (SFN)

Diabetes mellitus is a chronic disease characterized by insulin insufficiency (type 1) or insulin resistance (type 2, typically caused by obesity and metabolic syndrome), which in turn affects whole body energy homeostasis. Diabetes has the highest incidence rate of any metabolic disease in the world^4^, and is associated with numerous co-morbidities and increased mortality. The International Diabetes Federation reports that over 463 million individuals have diabetes, and it is estimated that 50% or more^5,6^ of patients with type 2 diabetes will develop peripheral neuropathy at some stage of the disease, while the range for patients with pre-diabetes is wider at 2%-77%. The majority of studies show a PN prevalence greater or equal to 10%,^7^ making it the most common complication of diabetes^8^. Diabetes is now the leading cause of PN overall, closely followed by aging/idiopathic. Since diabetes risk also increases with age, many diabetic cases are also age-related PN. More recently, COVID-19-associated Long COVID has become a new cause of PN^9^.

Over time it is thought that chronic hyperglycemia with diabetes causes damage to the capillaries and peripheral nerves of the extremities leading to DPN^10^. Other leading hypotheses implicate lipotoxicity instead of glucotoxicity, or a degenerative outcome from chronic inflammation^11^. Unlike nerves of the central nervous system (brain), peripheral nerves can regenerate to a certain extent, such as following nerve injury. Wallerian degeneration is a well-studied process of nerve repair following injuries like nerve crush, and involves Schwann cells and macrophages – however, this regenerative cellular cascade does not occur in cases of PN. Regardless of the initial trigger of the PN, once initial damage to the nerve begins, the initial response of the PNS is a regenerative one. During peripheral nerve regeneration a phenomenon known as ‘sprouting’ occurs, where the injured axons of peripheral nerves begin forming new branches^12^. In DPN, where the insult to nerves is continuous, this sprouting process can be prolonged and affect uninjured nerves, thereby contributing to increased pain sensation^12^. This often causes individuals suffering from early stages of PN to go through a period of hypersensitivity of the peripheral nerves, which can be painful. Following this hypersensitive period, progressive neural damage ultimately causes nerves to die back proximally in a length dependent manner (longer axons in distal extremities first), and from the skin surface down to deeper tissue layers, including, as we have demonstrated,^13–15^ in subcutaneous white adipose tissue (scWAT) and muscle. Small fiber nerve death can also progress to large fiber neuropathy, making PN an overall progressive and degenerative condition. Although peripheral nerves are capable of regeneration, something about the pathophysiology of PN blunts the ability of tissues to recover proper innervation spontaneously, and the ‘sprouting’ process is not akin to Wallerian degeneration with nerve injury. Particularly relevant in DPN, the neuropathy of metabolically important tissues such as muscle and adipose may further exacerbate the disease by blunting neural communication with metabolic command centers in the brain, such as the hypothalamus. Adipose tissue nerves, both the afferent sensory nerves and efferent sympathetic nerves, are known to be essential for lipolysis, thermogenesis, and other essential metabolic functions, and thus loss of this adipose nerve supply can have detrimental effects on metabolic control^13–15^.

Small fiber neuropathy (SFN) is the result of damage to both myelinated and unmyelinated small diameter peripheral nerves that innervate tissues and organs, as branches from larger nerve bundles. In humans, this includes Aδ- and C-fibers, which are considered small nerve fibers^16^. Aδ-fibers are 1-6 μm in diameter and myelinated, which accounts for their faster conduction velocity (4 to 36 m/s), as compared to the smaller (0.2-1.5 μm in diameter) unmyelinated C-fibers with a conduction velocity of 0.4 to 2.8 m/s^17,18^. In SFN, small somatic and autonomic fibers can be affected, which control thermal and pain perceptions, as well as autonomic and enteric functions^16^. As a result, patients with SFN can present with either autonomic or somatic symptoms, or a combination. Symptoms of small fiber neuropathy can include allodynia, burning, decreased thermal sensation, hyperesthesia, paresthesia, numbness in the extremities, restless leg syndrome, dry eyes and mouth, abnormal sweating, bladder control issues, gastric issues, skin discoloration, and cardiac symptoms such as syncope, palpitations, and orthostatic hypotension. As a disease, the pathology of small fiber neuropathy is poorly understood, but this is the type of neuropathy that is prominently observed in the leading PN etiologies: diabetes, aging, chemotherapy-induced, and now Long COVID^9^. SFN can also arise as a result of a variety of other diseases such as autoimmune disorders including sarcoidosis, paraproteinemia, and paraneoplastic syndrome.

### Limitations of Current Diagnostic Methods

Measurements of nerve electrical activity through functional, quantitative assays is possible for large fiber nerves (such as the sural or sciatic nerves) by techniques including nerve conduction velocity (NCV) or electromyography (EMG). While these methods are available in large medical centers as a diagnostic approach, similar methods for measuring functional changes to nerve electrical activity of the small diameter axons that innervate peripheral tissues and organs (those implicated in SFN like DPN) are not yet available. Therefore, the development of a simple diagnostic devise for SFN could expand an accurate, speedy, and cost-effective diagnosis across numerous clinical settings. Combinations of behavioral testing and patient questionnaires (qualitative data, rely on patient self-reporting), skin biopsies (fraught with high false negative and false positive rates and do not reflect nerve functional state), and NCV/EMG recordings (only capturing activity of large fiber nerves) are typically utilized for a diagnosis currently. Common problems that can be traced back to each of these testing methods include that diagnosing neuropathy is often a painful, invasive, and/or costly process that requires a medical specialist with unique expertise and equipment. Additionally, these approaches do not provide any quantitative, functional data on *small fiber nerves*, such as those innervating skin and subdermal tissues, and often result in results that are not definitive.

Skin biopsies, the gold standard diagnostic in SFN, are particularly fraught with high false negative and false positive rates since they are not indicative of nerve function or activity, and only a handful of labs in the country provide reliable analyses through immunostaining and quantification of intra-epidermal nerve fiber density (IENFD) from a skin punch biopsy sample. Furthermore, in SFN the pathophysiological changes in nerve function often precede anatomical changes such as those assessed by skin biopsy. Therefore, the lack of specific *functional* testing for SFN can result in unnecessarily long and arduous periods of diagnostic testing for both patients and clinicians, all while the disease is progressing and resulting in more damage to peripheral nerves. For less invasive yet still quantitative methods of SFN diagnosis, there is quantitative sensory testing (QST), or the quantitative sudomotor axon reflex test (QSART), which also have considerable limitations. QST is a well-defined sensitive method to examine thermal and mechanical sensory function, and as such is limited to assessing the function of large somatosensory Aβ-fibers and small sensory nerve fibers (ALJ- and C-fibers)^19^. On the other hand, QSART can only evaluate sympathetic cholinergic sudomotor function^20^. In conclusion, a measure of nerve function from all nerve types contributing to small fiber innervation of a tissue would greatly benefit diagnostic capabilities especially when used with current standards diagnostic testing, leading to improved patient outcomes.

### The New Detecting Early Neuropathy (DEN) Medical Device for Functional Assessment of Transdermal Small Fiber Nerve Activity

To provide a means of functionally testing for SFN and DPN by recording nerve electrical activity of all fibers innervating skin and subdermal tissues in a quantitative, quick, and sensitive manner, we developed a new medical diagnostic device. The Detecting Early Neuropathy (DEN) device is a needle electrode array consisting of 9 needle electrodes arranged in a 3×3 grid, with each needle connected to a copper trace on a single printed circuit board (PCB) that terminates at a cable connector. The DEN device needles are inserted into the skin, and penetrate to various subcutaneous tissue depths (typically 2-4mm), in order to measure nerve electrical activity over time, including in response to various stimuli like muscle flex or cold temperature. The recordings captured are needle pair differentials, providing a trace of all extracellular electrical activity and volume conducted field potentials, which are the electrical spread of signals generated by spatiotemporal integration of extracellular currents produced by neuronal activity ^21,22^ measured by those needles. Each needle pair is hooked to a single amplifier, which can also filter out room electrical noise. The purpose of this study was to validate and optimize the DEN device in a mouse model of DPN, and to determine if the DEN device can record from small fiber nerves in healthy human as a proof of concept for future clinical studies.

## Materials and Methods

### Construction of DEN Device Prototype for Mouse

The construction of the DEN device began with a prototype for assessing optimal spacing of the array needle electrodes. Plastic spacers were incorporated to allow varying penetration depth into the skin. The printed circuit board, or PCB, consists of 9 positions/holes arranged in a 3×3 grid, with each position attached to one needle electrode and a copper trace within the PCB, connecting to a single output pin of an edge connector (Fig. 1A-E). The experimental recording setup contained 4 amplifiers, from which each amplifier recorded from two needle outputs, one (+) and one (-) (Fig. 1F). Four different needle array designs were used, each with varying needle depths, in order to determine the optimal depth at which to record activity within the scWAT depot, as well as validate the functionality of the device during the recording process. After a preferential depth of penetration was determined, a single style of array was chosen to use for the duration of the high fat diet cohort recordings. The DEN platform is patent pending: international application number: PCT/US2021/031337, US application publication number US20230233822A1.

**Figure 1.**
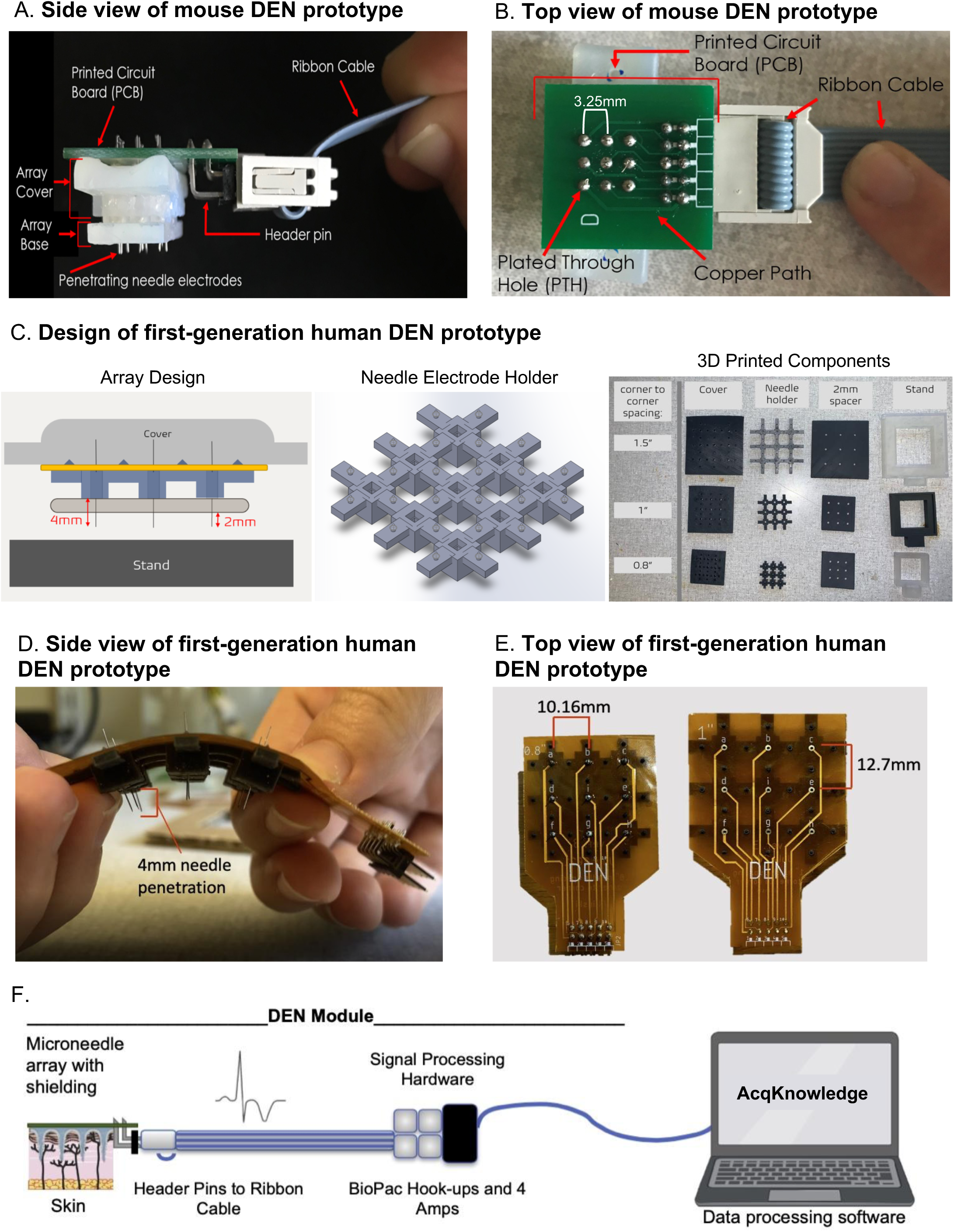
DEN Prototypes. Second generation mouse prototype: (A) side view, and (B) top view. First generation human prototypes: (C) Left: Array Design. Blue component is the needle electrode holder, yellow component is the flexible PCB, and tan component is the 2mm spacer. Middle: CAD design of needle electrode holder, for 1” array. Right: 3D Printed components of Prototype. Assembled human prototype (D) side view, and (E) top view. (F) Schematic of microneedle array with header pins attached to signal processing hardware and BIOPAC hookups.

### Animals and Dietary Intervention for Mouse DPN Model

Following validation of device nerve electrophysiology in cricket leg and limulus nerve, this study utilized weight-matched and age-matched male C57BL/6J (Jackson Laboratory, Bar Harbor, ME; stock number 000664) mice obtained from The Jackson Laboratory. Animals were housed with 4 per cage, providing for socialization in a temperature and humidity-controlled environment with a 12/12 h light/dark cycle. Access to food and water was maintained ad libitum and cages were changed daily. Following all recording sessions, animals were euthanized using CO_2_ followed by cervical dislocation. To achieve a diabetic state known to induce functional neuropathy, adult male C57BL/6J mice were fed 58% high fat diet from (Research Diets Cat #12330). Body weight was recorded weekly, and adiposity was assessed at the end of the study by weighing intact subcutaneous adipose depots after surgical removal. Mice were evaluated via electrophysiological flank skin/subdermal recordings in groups (N=4) at 2 weeks, 6 weeks, 10 weeks, 16 weeks, and 25 weeks after the start of the dietary intervention. Additionally, healthy control (chow-fed) mice (N=4) were evaluated at the same time points with flank skin/subdermal nerve recordings comparing the high fat diet mice against healthy control mice.

### Measurement of tissue nerve activity with DEN device- MOUSE

Mice were administered 5% isoflurane for 3-5 minutes in a takedown chamber, then continuously administered 1.5 - 2.0% isoflurane through a fitted nose cone for the duration of the recording. Complete recording sessions typically lasted no longer than 20 minutes per mouse, with at total time under anesthesia not exceeding 45 minutes per animal. Skin directly above the inguinal scWAT (∼1×1 inches) was shaved of fur using a razor blade to allow for better needle array penetration through skin. The mice were then placed inside a faraday cage, which redistributed charges from the exterior electric field in the recording room within a copper wire mesh, in turn preventing most exterior signal noise from interfering in the raw data. The DEN device needles were inserted into the shaved portion of the skin, directly over the subiliac lymph node, a region of known high density innervation^13^, then reinserted ∼1mm adjacent to the original penetration site twice, for a total of three penetration locations on each mouse during each recording session (Fig. 3A). All representative traces of raw data were taken within the first minute of recording from Position 1, Time 1. Following DEN insertion, the device was maintained in place by loosely tying the holding edge of the DEN device around the animal with plumber’s tape (Fig. 3B) and a 5-minute recording of nerve electrical activity was begun using a 4 channel, BIOPAC MP160 system. Amplifier settings were adjusted to 1000x gain, and a 0.5Hz high pass filter (HPF) and 3kHz low pass filter (LPF). Unless otherwise noted, mouse recordings were taken at 20kHz sampling rate. All mice were euthanized via cervical dislocation at the conclusion of each session.

### Measurement of tissue nerve activity with DEN device- HUMAN

For all human recordings, the DEN device was inserted into the lateral human calf above the ankle bone. Prior to insertion, the skin was disinfected with 70% ethanol and left to air dry. The subject’s leg was placed inside a Faraday cage and a ground was connected from the Faraday cage to the subject via an EKG surface skin electrode placed approximately 6 inches rostral of the DEN device. Following DEN needle electrode insertion, the device was maintained in place using self-adhesive athletic tape (ace bandage) wrapped around the calf and DEN (Fig. 6C). The Faraday cage was compared to wrapping the ace bandage with foil to insulate from room electrical noise, and it was determined that the cage was not needed. The DEN device was also disinfected with 70% ethanol and left to completely dry before insertion. Recording of nerve electrical activity was begun using the BIOPAC MP160 system. The BIOPAC recording system consisted of 4 individual amplifiers, each having two inputs. All 4 differential amplifier inputs were connected (each with a + and - input) to a 10-pin flat ribbon cable which was connected to the array via the PCB edge connector. The 9th input into the cable could be used as the ground reference, and the 10th input (located centrally on the array) was left open (to be used as a stimulation electrode if desired). For all recordings (unless noted otherwise), amplifier settings were adjusted to 1000x gain, 100 Hz high pass filter (HPF) and 3kHz low pass filter (LPF). Each amplifier was grounded to aluminum foil that was electrically connected to a Faraday cage. The raw data from this experiment was analyzed manually using the BIOPAC AcqKnowlege software. All recordings were taken at 40kHz sampling rate, allowing for the potential visualization of single action potentials. Due to the location of DEN insertion in the human calf, EKG signal was not factored into the rate and duration of patterned neural signaling observed with the BIOPAC software.

### Recording with EKG

When recording using DEN in conjunction with EKG equipment, a secondary area of fur was shaved over the chest of the mouse in addition to the typical area of the flank. A primary adhesive electrode was placed over the shaved area of the chest of the mouse. A secondary adhesive electrode was also applied to the hind paw. Both electrodes were connected to a single BIOPAC amplifier, allowing three channels to receive neural activity input from DEN and one channel to receive EKG. The DEN array was inserted into the flank of the animal as described in normal recording process, and both activity types were compared.

### Skin intra-epidermal nerve fiber density (IENFD) quantification

Glabrous hind paw skin and hairy flank skin (above the inguinal scWAT depot) was excised and fixed in 2% Zamboni’s fixative (Newcomer Supply # 1459A) for 2hrs at room temperature, transferred to 30% sucrose in 1XPBS overnight (until tissues sink), and embedded in OCT (Tissue-Tek cat #4583 Miles, Inc.) with orientation noted. Tissues were sectioned at 25 μm onto glass slides. Tissues were placed in 0.03% Typogen Black for 20 min at room temperature to quench autofluorescence, then rinsed in 1XPBS 3x 5min or until rinse ran clear. Tissue sections were blocked in 200 μL blocking solution (1X PBS/0.3% Triton X-100/5% BSA) at room temperature for 1 hr and then incubated in primary antibody (Proteintech rabbit anti-PGP9.5 (14730-1-ap), 1:1000) for 1 hr at room temperature and then moved to 4°C for incubation overnight. Sections were rinsed 3 x 1hr in 1XPBS and incubated with secondary antibody (AlexaFluor Plus 594 highly cross absorbed, 1:1000 (Molecular Probes)) for 1 hr at room temperature and then moved to 4°C for incubation overnight. Sections were rinsed 3×1hr with 1XPBS and then incubated in 100 ng/mL DAPI solution for 15 min. A final 3 x 10 minutes in distilled H_2_O rinse is performed and slides were then mounted with a drop of mounting fluid (Prolong Gold, Thermofisher #P36931) and a No. 1.5 glass coverslip for imaging on a Leica Stellaris 5 confocal microscope. Quantification of imaging was performed in Fiji/Image J. The pixel/μm ratio was set for all measurements as 4.5539 pixels/μm. The boundary of the region of interest (ROI) was created by outlining the entire epidermal layer on the 40x image. A measurement of the total ROI area was taken. The PGP9.5 channel was isolated and a consistent histogram criterion for thresholding was selected as ‘internodes’ across all images. The area of the thresholded nerves expressing PGP9.5 was measured, and the relative nerve fiber density was calculated: thresholded nerve area / total ROI area.

### BIOPAC Manual Data Analysis

The BIOPAC recording system consisted of 4 single amplifiers each containing two inputs. For the DEN needle array, all 4 differential amplifier inputs were connected (each with a + and - input) to a 10-pin flat cable for direct connection to the array. The 9th and the 10th input (located centrally on the array) was left open. The stimulation recording settings were designed for recording compound action potentials. For basal recordings, the amplifier settings were adjusted to 1000x gain, 0.5Hz high pass filter (HPF) and 3kHz low pass filter (LPF). For stimulation recordings, the amplifier settings were adjusted to 200x gain, 0.5Hz HPF, and 3kHz LPF. The raw data from the measurement of tissue nerve activity with the DEN device for mouse experiments was analyzed manually using BIOPAC AcqKnowlege software. Time periods were identified in which the first, second, and third recording locations occurred. An initial scan through the recording functioned to identify commonly occurring patterns of signals that existed in the raw data. Then, three small time (∼3-8 sec) intervals within the zone recording were chosen at random to analyze in depth. If familiar patterns of activity were observed within the chosen interval, the amplifier(s) on which the signal occurred as well as the size and duration of the signal were both recorded for reference. For manual analysis, recorded nerve traces were reviewed and action potentials (APs) were characterized by size and type according to predetermined parameters. Small APs were defined as signals just above the baseline noise to < 3x S/N ratio. Medium APs were defined as signals >3x S/N but <6x S/N. Large APs were defined as signals > 6x the S/N ratio, and very large APs were defined as >12 S/N ratio. Patterns of APs were characterized as a train (constant AP firing at specific rate for more than 2 msec), burst (burst of >1 APs for less than 2msec) or haphazard (single spike not part of a train or burst.

### BIOPAC Semi-automated analysis

Data analysis was performed with BIOPAC AcqKnowledge software. The spikes subject to investigation all had local voltage maxima and minima that could be defined in a cycle using the Find Cycle Peak Detector. To do this, event markers were placed at instances where a local maximum in voltage was above, or a local minimum below, a threshold that was defined based on the background noise. Using the Find Cycle Peak Detector, cycles were identified with maxima being the start events and minima being the end events, or vice versa if minima preceded maxima. Data generated and exported to excel from the Find Cycle Peak Detector included time of start event, change in time (Δt) between start and end event, maximum voltage in the cycle (M), minimum voltage in the cycle (m), and amplitude (A = M - m). By manual processing in excel, spikes defined as cycles were excluded if parameters were not physiologically relevant, i.e., if Δt > 3 msec and if 20 mV < A < 60 mV was not true. After exclusion of physiologically irrelevant spikes, change in time between any given starting event and the subsequent starting event (ΔT), otherwise known as the interspike interval, was calculated. An approximation for instantaneous rate of spikes was calculated using the formula 1/ΔT, and instantaneous rates that were physiologically irrelevant were excluded from analysis. To make comparisons between individual mice fed chow diet (N = 4) and 2, 6, 10, 16, or 25 wk HFD (each group N=4), instantaneous rates of spikes were averaged in intervals of 10 sec up until 350 sec. Only data in the first 350 sec of recordings were used to minimize the effect of anesthesia on comparisons between animals. For multi-class classification data were analyzed using a proprietary algorithm owned by Neuright, Inc.

### Statistical Analysis

Mice were age and body weight matched and randomized to treatment groups. All plotted data represent mean +/-SEM. Statistical calculations were carried out in Excel or GraphPad Prism software (La Jolla, CA, USA), utilizing ANOVA, or Student’s T-test as indications of significance (specified in Figure legends). Error bars are SEMs. For all figures, *p < 0.05, **p < 0.01, ***p < 0.001, ****p < 0.0001.

### Randomization, Blinding, Replicates, Inclusion/Exclusion Criteria

For animal studies, mice were age-matched and randomly assigned to a dietary group. A total of 4 biological replicates were evaluated in nerve activity recordings for each dietary group. Experimenters were not blinded to animals group assignment during nerve recording sessions; but experimenters were blinded for the assessment and analysis of nerve recording data. All measured data was included in analyses and reported.

## Results

There were four major goals of this project, which included: 1) validating the functionality of the DEN device to detect nerve electrical activity, through testing in invertebrates and control mice, 2) determining the optimal tissue depth, needle fabrication, and anatomical placement of the microneedle array for mouse pre-clinical studies, 3) comparing control recording data with a time-course of diet-induced DPN progression in mice, against other measures of DPN/SFN including IENFD, and 4) fabricating a human prototype for testing DEN recordings in healthy human.

### Device designs for mouse model and human use

The fundamental components of the DEN were present in both mouse and human device prototypes, while major differences between species designs included device size and flexibility of the array due to printed circuit board (PCB) choice (Fig. 1). Figure 1A-B illustrates the DEN as optimized for use in our mouse models. Each array had three parts: base, cover, and stand. The base and covers had nine cutouts in a 3×3 array, that allowed for 4 needle pair differential recordings, leaving 1 needle in the center of the array available for electrical stimulation, if desired in the future. Custom needle electrodes were inserted into the base, which were used to collect nerve activity data from subjects. The second-generation mouse prototypes used in this study had needle electrode centers that were spaced 3.25mm apart. Arrays were designed to have penetration depths up to 2.5mm to target mouse skin down to the scWAT layer. Arrays either had the same penetration level for all nine needle electrodes in the array or had varying depths across the array. The base, cover, and stand were 3D printed with a Formlabs Form 2 printer with the “standard” resin, which is hard and sturdy. The needle electrodes were housed in a base needle electrode holder, where the needle electrodes extended out the bottom of the holder. When the holder was placed onto a subject, the needle electrodes penetrated through the skin to make the transdermal recordings. A PCB connected to a ribbon cable was used to electrically connect the array to a recording and amplification system (commercially made by BioPac).

Following validation in the mouse models, a first-generation human prototype was constructed and tested in two healthy volunteers. The PCB for the mouse prototypes (Fig. 1A-B) is a glass reinforced epoxy (green), and in the human prototypes is a flexible polyimide laminate (Fig. 1D-E). The major differences between the mouse and human prototypes were: 1) the increased spacing between the needle electrodes, 2) penetration depth of the needle electrodes into the intended subjects (up to 4mm penetration depth in human), and 2) the rigidity of the prototypes, where flexibility of prototypes was quantified as having a bend radius less than 1.5” to conform to the curvature of the human body, to allow for use over different skin surfaces.

The DEN prototypes designed for human use contained the same components as mouse, including PCB, needle electrodes, 3D printed needle base, cover, stand, and cable with connector. A diagram to visualize the assembly of the components into the prototype is shown in Fig. 1C, left panel). Needle electrodes are inserted into the needle electrode base (Fig. 1C, middle panel), which is designed to allow 4mm of the needle tips to extend out of the base. The human needle electrode base was designed to maximize flexibility in the array while maintaining sufficient support for each individual needle electrode, so that the array could conform to the human skin surface while keeping each needle in proper alignment for orthogonal penetration. The final design featured a 3×3 array of rigid individual needle holders connected by flexible arms, and mechanical attachments to connect the needle electrode holder to the flexible PCB. The final design of the individual needle holders were 7 mm x 7 mm x 4.75 mm rectangular blocks with 2.3 mm x 2.3 mm openings at the center, into which the custom needle electrodes were pressed and held firmly in place. The connectors between the individual needle holders varied in size depending on the overall size (and spacing) of the needle array. Three different prototype sizes were developed for the project, two of which are shown in Fig. 1E. The smallest had the distance between needle centers 10.16mm apart. The middle prototype had needle centers 12.7mm apart, and the largest had centers 19.05mm apart. The sizes reflected a wide range to test for optimal spacing for assessing neural health. Fig. 1C (middle panel) shows the final needle electrode holder design. These are examples of the near-infinite iterations of array sizes and needle spacing that are possible with these custom array designs, allowing us in future to optimize design for different skin surfaces such as torso, arm, or leg.

A 2mm optional spacer can be added below the needle electrode base, to reduce needle penetration depth if desired (Fig. 1C, right panel), and a cover for the array was created for application safety and to assist with insertion of the prototype by the operator. Because the needle electrodes extended above and below the needle electrode base (for other functions of this device as a theragnostic that can sample interstitial fluid for biomarker discovery and testing, and to deliver treatments as we have done previously^23^, a cover with height greater than 5mm was needed to ensure an administrator would not be at risk for a needlestick injury, and to apply an evenly distributed force to all of the needles in the array during insertion into a patient’s skin (and thus avoiding any needle damage/bending during the process). Stands were designed to support an assembled prototype, which lifted the array by 10mm, to ensure needle electrodes do not come into contact with the surface of the stand. The base, spacer, cover, and stand were 3D printed with Formlabs Form 2 printer with the “Flexible V2” resin. The flexible PCB is much thinner than traditional PCBs and is specially designed to withstand bending. The flexible PCB was placed on top of the needle electrode base and electrically connected to the needles (Fig. 1D). For both mouse and human recordings, the DEN was connected via ribbon cable to signal processing hardware and the BIOPAC amplifiers (settings described in Methods section), see schematic in Fig. 1F.

### Validation of the DEN Device

Prior to vertebrate studies, experiments were performed in invertebrates to optimize device parameters and to confirm that nerve electrical activity was being detected and recorded as expected, by users who typically record from nerves of invertebrates. For confirmation of signal parameters, we compared our DEN needle electrodes with commercial Natus electrodes (Natus Medical Incorporated) in cricket leg recordings. Natus electrodes are disposable electrodes often used in nerve conduction velocity (NCV) diagnostic tests in mouse and human^24^. Similar to the custom DEN needle electrode, the Natus electrode is made of stainless steel. In these experiments, a single leg was removed from two anesthetized crickets. In each leg, an electrode, either DEN or Natus, was placed in the coxa, and a ground was placed into the tarsus. Following electrode insertion, recording was started. A mechanical stimulation was supplied to the cricket leg through a puff of air. This stimulation caused compound action potentials in the leg, which were recorded and displayed using the BIOPAC Acqknowledge software (Suppl. Fig. S1A). Limulus nerve was used to determine optimal needle spacing for differential recordings within the DEN array across needle pairs connected to a single amplifier (Suppl. Fig. S1B).

Three additional experimental trials were performed in anesthetized mice for the purpose of validating the DEN electrodes against commercial Natus electrodes, as well as to test the functionality of the DEN second generation mouse device as a unit recording extracellular axon potentials in transdermal tissue. In all mouse validation studies, the Natus electrode provided stimulation (which in the end was deemed not necessary to record baseline transdermal recordings, since we saw random firing and spontaneous action potentials in the transdermal signals, without electrical stimulation).

Data were collected in mouse using both Natus and custom DEN needle electrodes. The first experiment used an electrical stimulation from the BIOPAC system into the mouse’s sciatic nerve and recorded the response from the right front paw. Since stimulation in this experiment was electrical and we are also recording electrical activity in the tissue, a stimulus artifact was expected to appear in the raw recordings. Stimulus artifacts appear as a sudden spike of voltage on the recording trace, followed by a return to the voltage values prior to stimulation, and ending with a plateau at the initial voltage amplitude prior to a stimulus response. Compound action potentials (CAPs) followed the stimulus artifact and typically appeared as signals with near-symmetry above and below the x-axis, with a smooth curve at the peak. CAPs were detected at a 50mV stimulation and 200x gain from all three recording differentials (Suppl. Fig. S1C top 3 panels).

We next recorded from mouse flank without electrical stimulation, and confirmed that the DEN electrodes could obtain nerve recordings from a live mouse, as shown by the signal generated (Suppl. Fig. S1C bottom panel). We consider these recordings as basal recordings of nerve function. The electrodes recorded differentials across two custom needle electrodes placed in the tail, with a ground in the front right paw of the mouse. These recordings without electrical stimulation also ensured that all connections, hardware, and software functioned properly. The large 0.5mm, 1.0mm, and 1.5mm penetrating array designs were used for these recordings.

The third experiment recorded from the mouse flank region using the intact DEN device, plus one custom needle electrode penetrating the paw, similar to the NCV diagnostic test done in humans. All differentials displayed a stimulus artifact at a 200x gain when a 50mV stimulation was supplied to the array (Suppl. Fig. S1D). It was determined that the differentials located closest to the stimulation show the greatest stimulus artifact. In addition to stimulus artifact responses, CAPs could be seen clearly in Channels 2 and 4 of the four differentials.

To examine if enhanced conductivity was necessary to improve recordings obtained from our coated stainless steel needles, a thin silver wire was placed inside the hollow needle electrodes of the DEN during recordings, and then the wire was removed while the DEN device recorded (see Fig. 3A for our positioning schematic, we tested 3 sites over time above the flank skin region). After anesthetization and insertion of the array, the silver wire was pulled from each single needle, and recording was stopped and re-started after the removal of each wire. The silver wires were removed according to the channel each needle corresponded to, with the wire removed in ascending channel order (Channels 1, 2, 3, then 4), in order to determine if the silver wires were essential for design functionality. Finally, the DEN device recorded with no silver wire through any of the needles for several minutes. The raw data recordings revealed no difference in the prevalence or rate of electrical activity recorded from the control mouse (Suppl. Fig. S1E) regardless of silver wire placement, and thus wires were not included for subsequent recordings.

### Neuropathy Time Course Study

In order to correlate nerve electrical signals measured by the DEN to commonly used assessments of SPN, we employed a mouse model of diabetic/pre-diabetic peripheral neuropathy that is induced by 58% high fat, high sugar diet feeding for at least 16 weeks, as previously validated by NCV,^25^ and also verified independently by our laboratory by von Frey in previous studies^26^. We employed this high fat neuropathy-inducing diet (HFND) over a time course before and after neuropathy initiation at 16wks of diet, to evaluate changes in peripheral nerve activity at pre-diabetic stages, as well as later neuropathic stages in this pre-clinical mouse model of DPN. A diagram of the diet time-course study design is in Fig. 2A. It is important to note that functional large fiber neuropathy has been seen in mice at 16 weeks after consuming this diet, but in previously published studies paw skin IENFD remained unchanged at this time point^25^. This further underscored the disconnect between IENFD and nerve activity/function, and the need for additional diagnostic measures beyond histological quantification of nerve ending terminals. On the other hand, studies have shown that at 36 weeks of high fat diet feeding a striking reduction in IENFD is evident in mouse paw skin^27^. Therefore, 2 week, 6 week, and 10 week mouse cohorts in this study were included to precede the point at which functional large fiber neuropathy was expected to occur (16 weeks), as previously demonstrated by NCV, while the 25 week point was included in order to evaluate post-neuropathy onset timepoint^25,27^. We anticipated that small fiber nerve activity measurements even at these early time points may show patterns different from lean (chow) controls, indicative of diet/obesity-induced neurodegeneration. In particular, the recording of any compensatory effects due to nerve sprouting prior to measurable functional neuropathy was a particular point of interest throughout this project, since it is not easily detected in clinical populations. As expected, body weight and scWAT weight increased the longer mice remained on HFND (Fig. 2B). We could not control for mechanical effects of a thicker scWAT layer under the skin that may have displaced needle penetration, but the array was always placed flush with the skin surface for recordings (and in later studies, we have validated SFN in a non-obese model, indicating subdermal adipose is not a confounder). As expected, there was a steady reduction in IENFD over time on the diet, with a significant decrease in nerve density at the 16-25 weeks HFND time point compared to 2 weeks on HFND (Fig. 2C). This fits with prior observations^27^.

**Figure 2.**
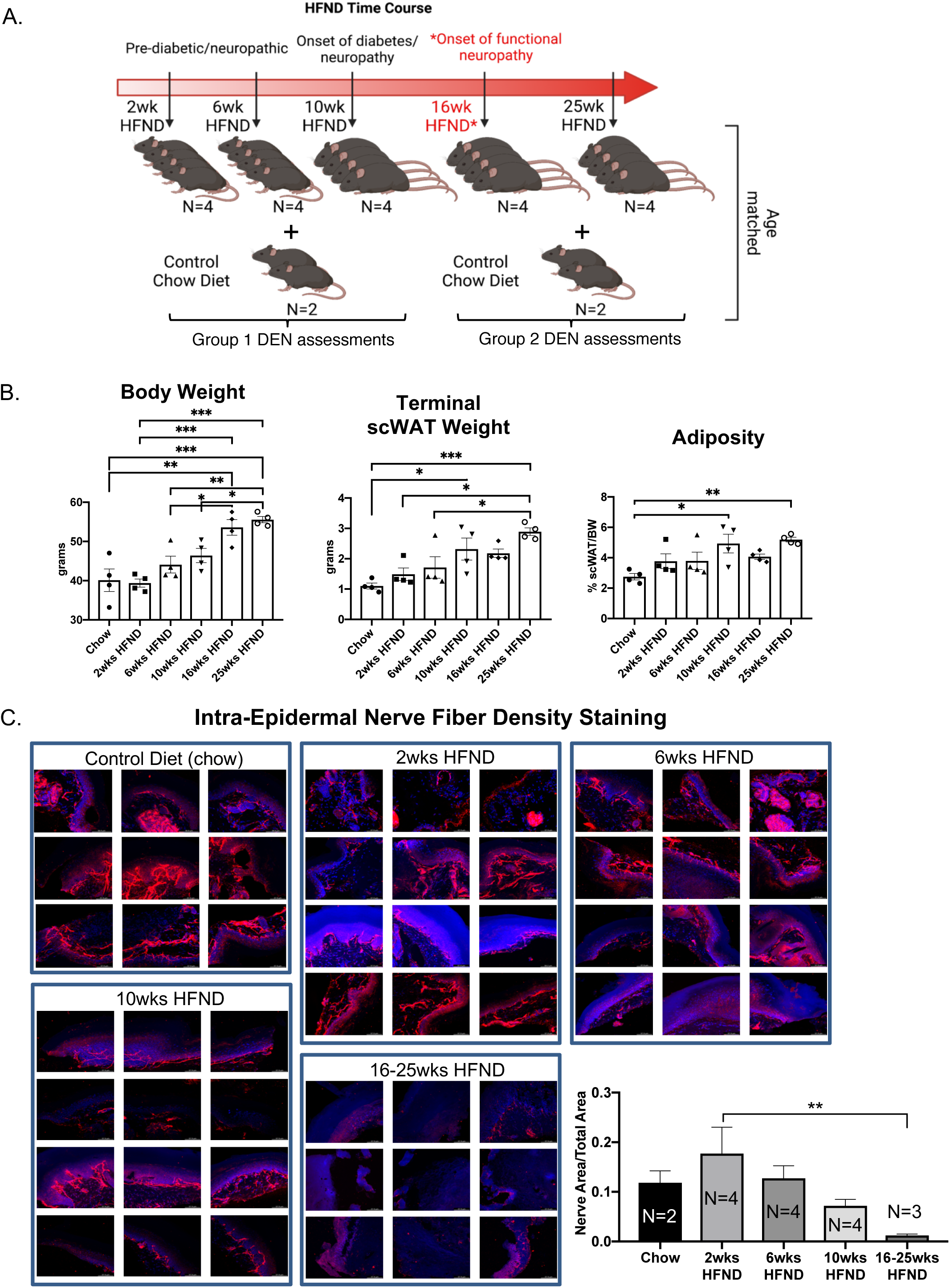
Study design for assessment of diet-induced neuropathy. (A) Visual schematic of study design: adult male C57BL/6J mice were placed on a 58% high fat neuropathy-inducing diet (HFND) for a time-course intervention. Critical physiological changes induced by HFND are denoted with respect to duration of HFND feeding. Assessments were performed at 2, 6, 10, 16, and 25 weeks after dietary intervention. *Neuropathy is evident by 16 weeks of HFND feeding. (B) Body weight of HFND and chow diet controls at time of assessment. Body and tissue weight data were analyzed with a one-way ANOVA and Tukey’s post hoc test. For B, N = 4 per group. Error bars are SEMs. (C) IENFD of glabrous hind paw skin showing PGP9.5 expression in the epidermal layer, results are representative of 3-4 images per animal.

Three major anatomical zones were used for recordings in mice using the DEN device. For Position 1, Time 1, the array was centered above the subinguinal lymph node (SiLN) region, since this area has been shown to have a dense innervation of nerve fibers in the inguinal scWAT ^13^. Position 2, Time 2 was located approximately 1mm rostrally to Position 1 Time 1, and Position 3, Time 3 was located approximately 1mm caudal to Position 1, Time 1 (Fig. 3A). Moving the needle array slightly throughout the recording sessions allowed for variation in the types of activity seen, since these needles pick up extracellular field potential signals from all nerve endings within their vicinity, and this also increased the chance that the array needles would become positioned optimally near a single axon, or bundle of nerve fibers, so we could observe different patterns of nerve activity. Prior to recordings, mice were anesthetized and stabilized for the insertion of the DEN array (Fig. 3B).

**Figure 3.**
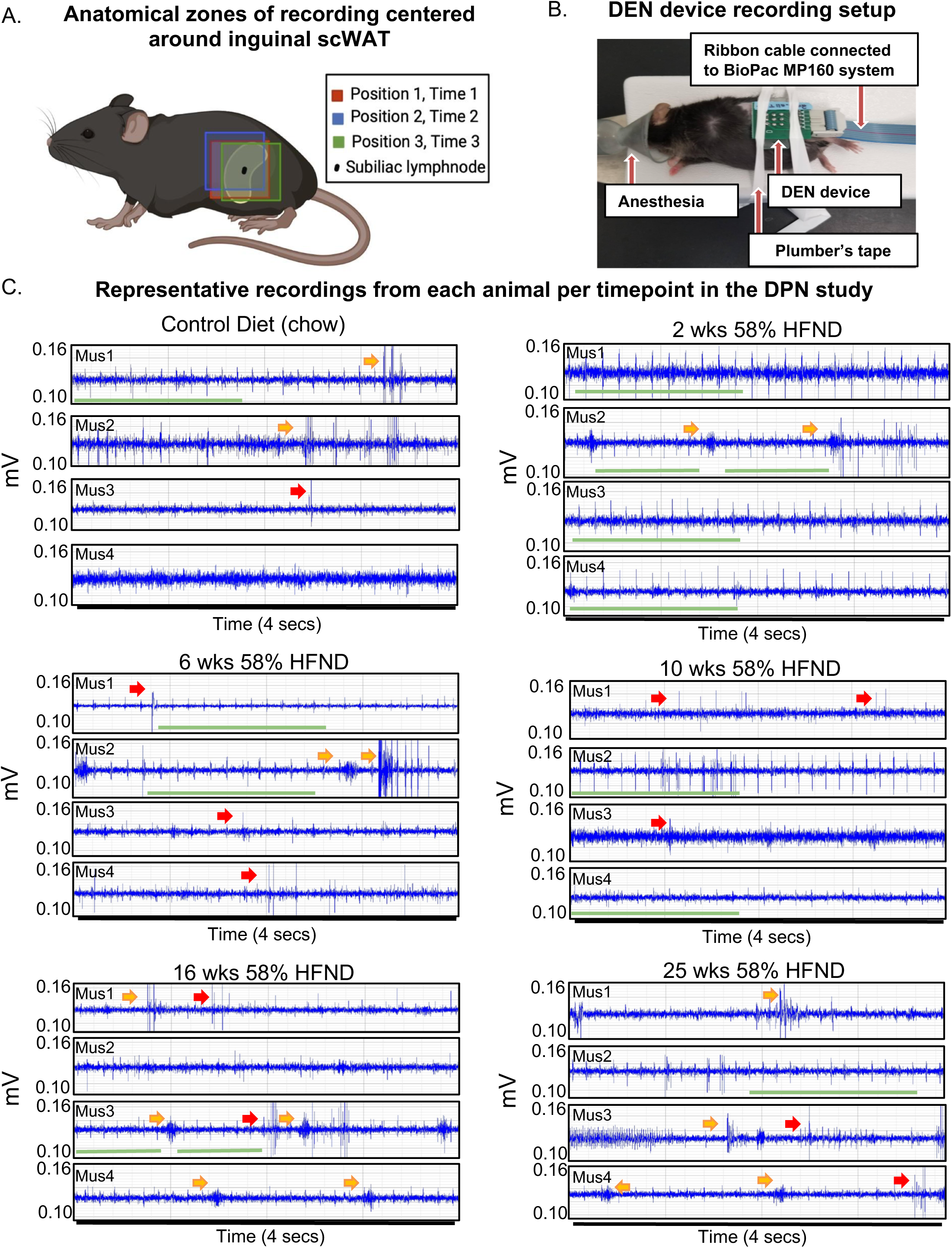
DEN placement and representative recordings from animals on HFND. (A) Visual schematic showing array insertion locations relative to the subinguinal lymph node. (B) Representative mouse from preclinical trials shown with array inserted over the left inguinal scWAT depot. (C) Representative electrophysiological traces from individual mice (mus1-mus4 per dietary group) used in the study. All representative traces were taken within the first minute after placement of array in Position 1; each trace recording represents a 4 second duration. Voltage range remained between 0.10-0.14 mV with amplitudes of signals often appearing at greater than 2:1 signal to noise ratio. Green bar denotes spike train, yellow arrow points to a burst, red arrow points to single spike.

The DEN recorded a variety of nerve activity signatures across the four needle pair differentials (i.e.: four amplifiers), including putative action potentials (PAPs) that were observed across all time points in the HFND longitudinal study. There were three main types of activity patterns identified, which occurred across all three recording zones in both control and HFND treated mice. The first type of activity was spike trains, defined as the repetitive and constant firing of PAPs over a set period of time (Fig. 3C, Suppl. Fig. S2A). This property allowed spike trains to be traced across the different time points of HFND. The second type of activity recorded was bursts, or the firing of many PAPs over a short period of time, typically within less than 1 second (Fig. 3C, Suppl. Fig. S2A). Finally, single PAPs were recorded as isolated events that appeared similar to neural action potentials (Fig. 3C, Suppl. Fig. S2A). Criteria used to identify PAPs included the presence of relative symmetry of the signal amplitude both above and below the x-axis, a signal duration of less than 0.5 milliseconds, and the non-synchronous presence of the signal on more than one recording channel. Later computational methods confirmed these manual observations, as described below.

Since time under anesthesia was a confounding variable that was also not present in human testing, we measured impacts from anesthesia separately (Suppl. Fig. S3A), and concluded that anesthetic impacts on the nervous system (such as the hypothermia that occurs under inhaled anesthesia), could also be captured by the DEN over time (Suppl. Fig. S3B). This was therefore controlled for in our final statistical and computational analyses.

### Spike Trains are Consistent with EKG Signals

By simultaneously recording with the DEN and traditional surface electrodes placed on the mouse for EKG recording, spike train patterns were able to be manually analyzed alongside known EKG signals using the BIOPAC equipment and software to determine if DEN signals included EKG signals in the mouse. Upon completion of this analysis, it became apparent that certain spike train signals were occurring at the same time signature and rate as the EKG signals in the mice as measured by surface electrode, likely due to how close the heart was to the flank recording site in the mouse (of note, this was not a confounding signal in future human recordings, however, since the device in the leg was not close to the human heart). Both the spike train and EKG signals occurred at the same rate and fluctuated in tandem throughout the recording session (Fig. 4A). Using a proprietary algorithm developed by Neuright, Inc. as part of the FOX-DEN software (fidelity operation transfer for DEN, or FOX-DEN platform), we were able to further process the DEN data through a 300Hz 10^th^ order High Pass Filter and analyze the EKG signal separately and optionally in our DEN recordings. We found that the EKG spike train produced only 5 spikes per second, while we measured a mean spike rate of 500 spikes per second from the DEN recording. Therefore, the EKG signal accounted for only 1% of the total mean spikes within the DEN recorded data and was not contributing to overall interpretation of PN. Interestingly, the overall rate and prevalence of the EKG spike train activity decreased significantly at only two specific time points of HFND feeding (2 weeks and 25 weeks of the HFND compared to chow controls) (Fig. 4B). At the 25 week time point, functional and histological PN is present in skin, adipose, and muscle, and the decrease in spike train/EKG activity is likely indicative of additional cardiac autonomic neuropathy (CAN), a known condition in obesity and diabetes that is also often underdiagnosed^28,29^. The changes observed at the 2 week time are also interesting, as leptin resistance is present in mice by 2 week of HFD feeding, and this may impact EKG signal ^30^.

**Figure 4.**
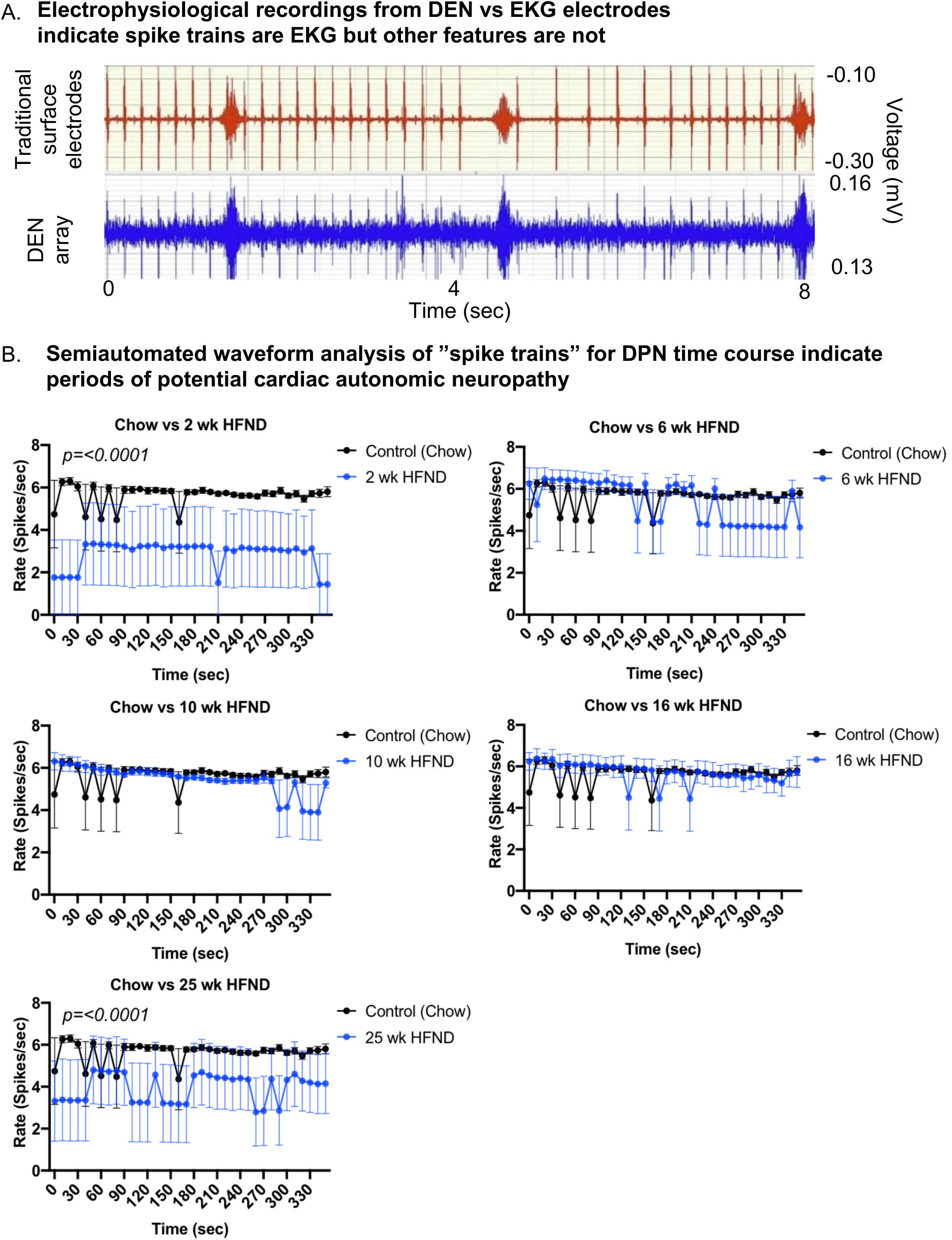
Comparison of DEN recordings to EKG signal. (A) EKG recordings using traditional skin surface electrodes (top channel, red) occurred in synchrony with spike train signals (bottom channel, blue) recorded from DEN positioned in murine flank. EKG skin electrode was placed over the shaved chest of the mouse and ground was attached at the front paw. (B) Semiautomated waveform analysis of “spike trains” for DPN time course. Two-way Ordinary ANOVA, for statistically significant findings Source of Variation is Diet. For all error bars are SEMs.

### Statistical learning-assisted modeling from mouse recordings using the DEN

The first 10 minutes of nerve activity in transdermal tissue was captured from each animal in the neuropathy time course study and was used to determine how statistically different nerve recordings were across the HFND time points, compared to control animal recordings. Clustering visualization using Principal Component Analysis (PCA) dimensionality reduction revealed that nerve recordings maintained discreet and separate clusters that were dependent on length of time on HFND (Fig. 5A). Each point represented the full DEN data associated with each mouse in a given HFND group. Data in the high dimensional feature space was rotated and projected into a 3D space for visualization by using PCA (Fig. 5A). Models were constructed and trained for all data using 5-Fold Cross Validation. Based on these discoveries, we constructed a multi-class (4-class) classification model. The features for Statistical Learning were obtained by splitting the 10 min data into 4 sec segments, and therefore only 4 sec of nerve recording data was needed in order to predict to which HFND group the nerve signature belonged. The multi-class classification model gave an average 5-Fold Cross Validation accuracy score of 91.73% with a standard deviation of 0.78%. Our model was able to classify mice in the 16wks HFND group with 96.94% accuracy (Fig. 5B). This matrix is comparing the actual HFND groups (vertical axis) with those predicted by our Statistical Learning mode (horizontal axis). For example, the upper left square is indicating that of all the Chow Control predictions made by our model, 91.75% were correct. In fact, of the neuropathy time course groups used in our model (chow, 6wks HFND, 10wks HFND, 16wks HFND) all predictions achieved an accuracy higher than 90% (Fig. 5B). This modeling indicates that our data collection method yields information that is correlated with nerve electrical activity signature and length of time on HFND, providing preliminary support for the clinical diagnostic relevance of our data.

**Figure 5.**
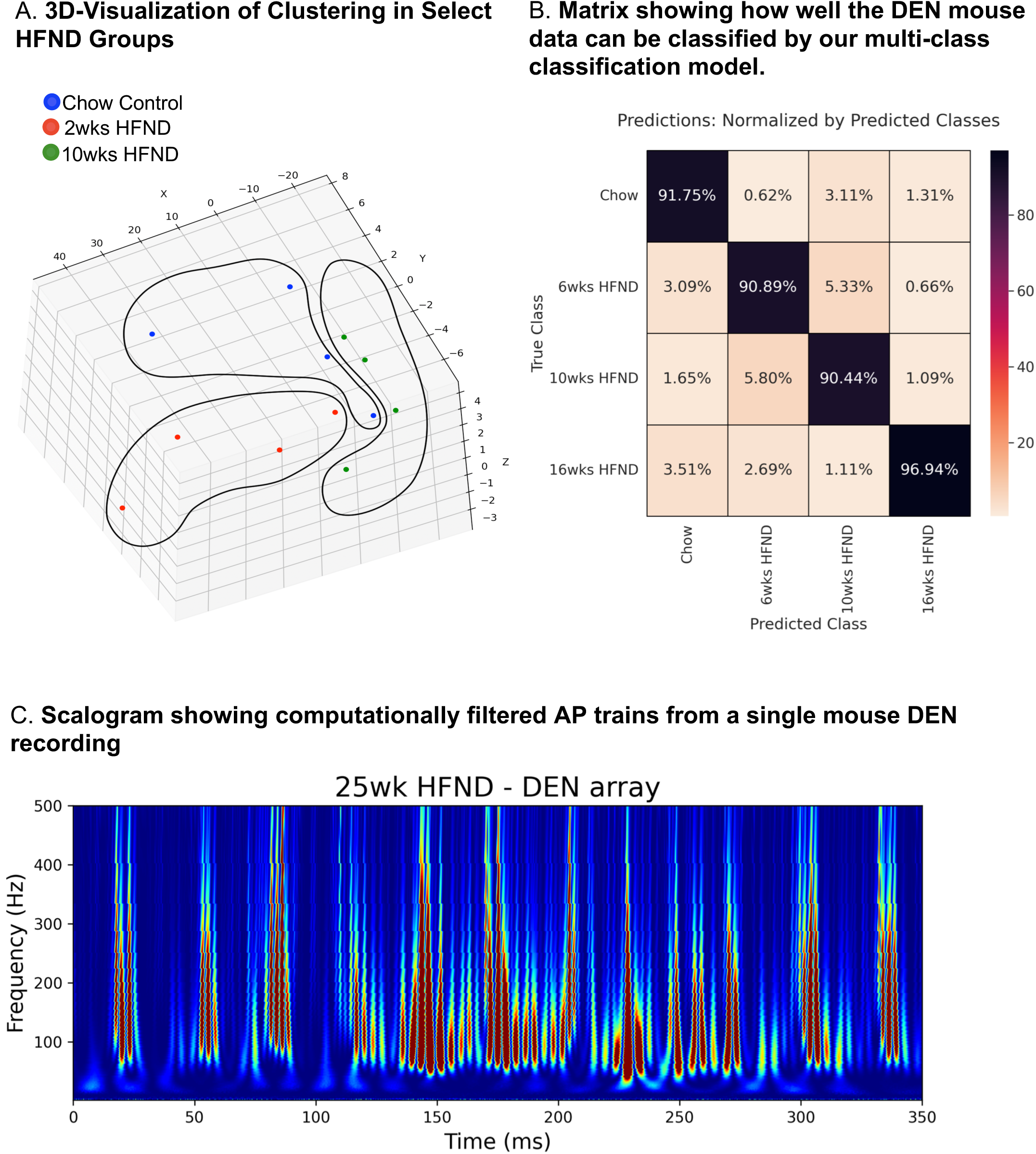
Statistical-learning-assisted modeling of DEN recordings in mice across a HFND. (A) Visualization of Clustering in Select HFND Groups. Each point represents 10 min of DEN data associated with each mouse in a given HFND group. Data in the high dimensional feature space was rotated and projected into a 3D space for visualization by using PCA (Principal Component Analysis). (B) Confusion Matrix showing how well the DEN mouse data can be classified by our multi-class classification model. This matrix is comparing the actual HFND groups (vertical axis-True Class) with those predicted by our Statistical Learning mode (horizontal axis-Predicted Class). For example, the upper left square is indicating that of all the Chow Control predictions made by our model, 91.75% were correct, and the lower left square represents the percentage of Chow Control predictions made by our model whose true group was 16wks HFND. (C) Custom FOX-DEN software was used to generate a scalogram using data obtained from a mouse DEN recording from the 25 wk HFND group, each peak is representative of a recorded action potential.

### Computed Action Potentials

We next used the Continuous Wavelet Transform (CWT) scalogram^31,32^ to visualize the signature of action potential trains present in a time series of extracellular recordings. To illustrate the signature, we first theoretically simulated (Suppl. Fig. S4A) a noise-free time series consisting of computed action potential trains (Suppl. Fig. S4B). We also computed the scalograms of data consisting of pure Gaussian noise (Suppl. Fig. S4C) and showed that these scalograms looked very different and did not contain the signature of computed AP trains exhibited by the theoretically computed AP train (Suppl. Fig. S4B). For the computed action potential, we used a simple theoretical simulation of an action potential *v(t)*, shown in (Suppl. Fig. S4A), (time *t* is in ms),

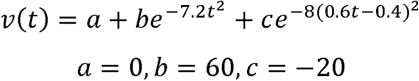

Note that by selecting a=0 we are subtracting the resting potential from the action potential. The results of our computational analysis are not critically dependent on the precise shape of the computed action potential. Any curve with a general shape similar to an action potential will yield similar results.

For a noise free AP train of N action potentials, v_train (t), we use:

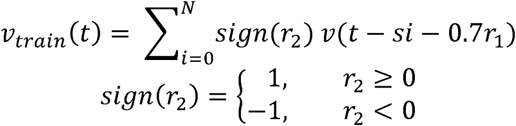

*r*_1_ and *r*_2_ are random numbers lying in the interval [-0.5,0.5]. New random numbers are generated at each summation. *s* determines the spacing between the action potentials. When *s* is small (i.e., *s*= 1) the action potentials overlap with positive or negative signs (i.e. the *sign*(*r*_2_) factor) to form compound action potentials. The 0.7*r*_1_ term, places the action potentials not exactly at periodic time intervals but offsets them by an amount 0.7*r*_1_. Note that *v_train_*(*t*) does not contain a noise term added-on to the signal as a sum. The random terms simply move the action potentials in time to simulate compound action potentials measured from various neurons in the neighborhood of the measuring needles. In Suppl. Fig. S4B, the contours of the scalogram of such a computed train were shown to exhibit the typical signature of a noise-free computed AP train. Each red contour corresponds to an action potential. The red contour starts from a body (thicker end of the red contour) at around 100 Hz and becomes thinner at higher frequencies. Since the action potentials partially overlap, the bodies of the red contours do not all lie at the same frequency, rather the bodies lie on a wavy curve as time increases at the bottom of the scalogram image. For comparison, we generated the scalogram of pure Gaussian noise (Suppl. Fig. S4C). Note that the contours of these scalograms do not exhibit the signature of an AP train. The bodies of the contours do not all start from roughly the same frequency, instead, they originate from randomly arbitrary (low and high frequencies) throughout the whole area of the scalogram.

Using this computational approach, we were able to show that the DEN recording from one of our 25wk HFND animals generated a scalogram of the same signature exhibited by the noise-free computed CAP train (Fig. 5C). This scalogram illustrates that the DEN data is mostly composed of CAP trains, authentic nerve electrical signatures, with low noise levels. Furthermore, as the DEN recordings are taken at basal conditions without any electrical stimulation applied, discrete physiological changes in nerve firing rates and patterns may be more easily discerned.

### Optimization of the DEN device for human studies

Upon completion of pre-clinical studies in diet-induced models of PN in mice, the DEN device was optimized and re-designed to be tested on the lower leg of two healthy human subjects. There are notable differences between murine skin and human skin (illustrated in Fig. 6A) which needed to be considered including: greater epidermal and dermal thickness of human skin, fewer hair follicles in human skin, and presence of eccrine sweat glands. Additionally, human skin is firmly attached to underlying tissue layers (mice have looser skin) and lacks the *Panniculus Carnosus*, a muscle layer present in mice and other mammals that is involved in skin contraction, wound healing, and skin adherence to underlying tissue ^33^. The spacer component was used with the DEN device to reduce the needle penetration into human skin from the maximum depth of 4mm to the minimum depth of 2mm for initial recordings (Fig. 1C, 6B). Since human skin thickness varies greatly depending on location, averaging from 1-4mm^34,35^, this design choice was made to ensure that we achieved nerve recording throughout the depth of human skin and underlying tissues. As the human subject testing was being done on the lower leg, only the 2mm needle length array was tested to avoid unnecessarily painful needle insertion. Indeed, the two participants reported minimal or no pain upon device insertion to the leg, and once inserted the device was barely felt. This 2mm depth of penetration allowed for a full skin (epidermal and dermal) recording while reaching the subcutaneous layer as lateral lower leg skin thickness is approximately 1.7mm thick^36^ (Fig. 6A). The device was inserted into the lateral calf approximately 4 inches above the ankle, and secured in place using self-adhesive athletic tape (Fig. 6C) with foil lining to serve as a Faraday cage (reducing room electrical noise contamination in the data). Participants were asked to remain still for 30 seconds, then move their foot and ankle at various points during the recording session to initiate the EMG signal in the data. These time points were marked in the trace and as metadata.

**Figure 6.**
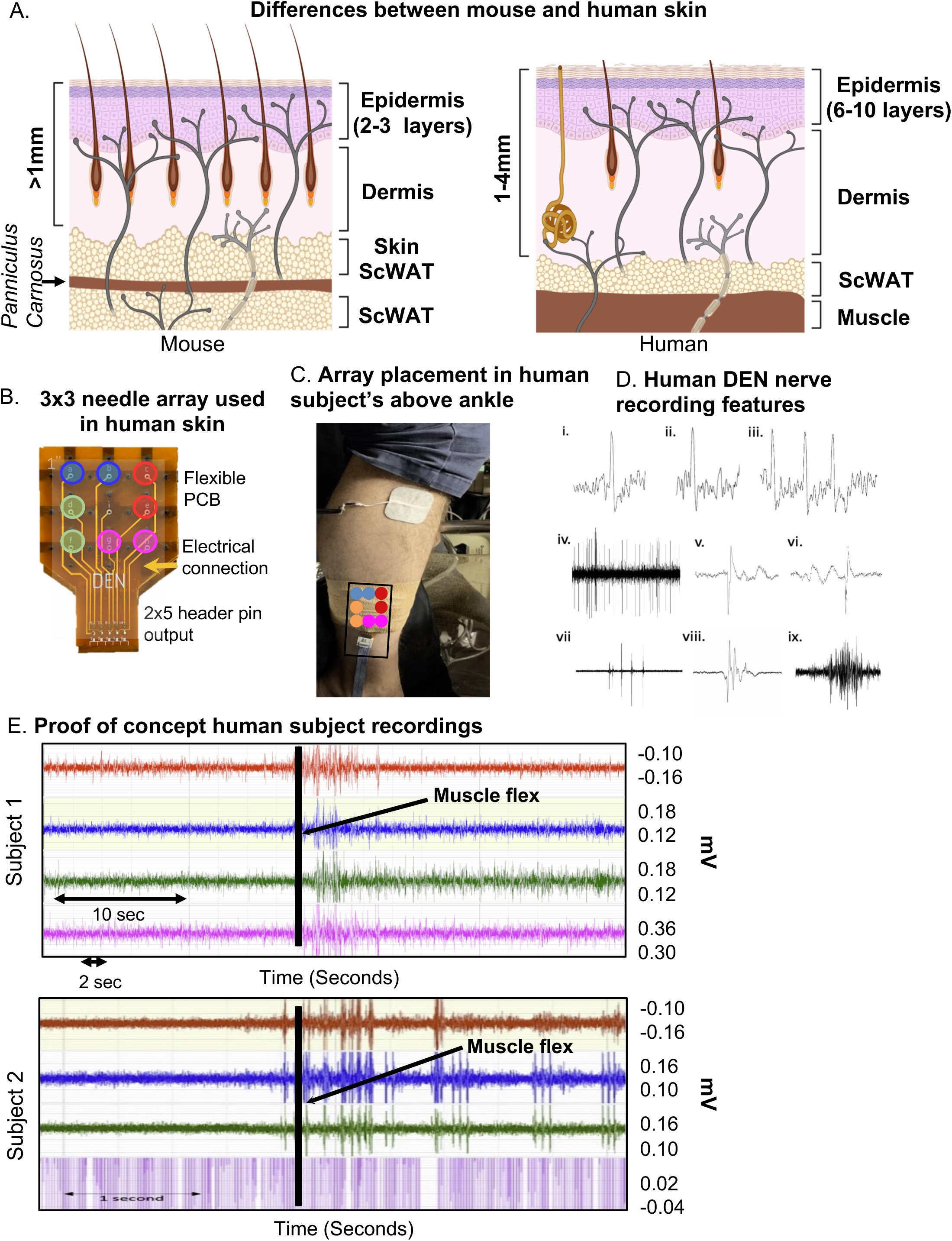
Optimization of DEN for human recordings. (A) Simplified comparison of skin layers between mouse (left) and human (right) showing differences in epidermal and dermal thickness, and underlying scWAT. (B) Human DEN prototype with 3×3 needle array and corresponding output connections to recording amplifier channels. (C) DEN inserted into lateral human calf. (D) Single Action Potentials (i-ii) and a short burst of three APs (iii) are recorded from a human subject. The APs are each about 30 mV in amplitude within a baseline noise of about 10 mV. Durations of traces are approximately 5, 4, and 11 msec long. A train of CAPs/CMAPs (iv) recorded from subject NE. 4 pattern bursts, two early on and two later, of CAPs/CMAPs occur within a train of regularly firing CAPs/CMAPs. Peak to trough amplitudes within this eight sec trace are approximately 180 mV. Two of the CAPs/CMAPs (v-vi) from near the middle of the train show time resolved (trace durations of about 8 msec each) details of these CAPs/CMAPs, along with slower waves of electrical activity before (vi) and after (v) the CAPs/CMAPs, recorded at 40kHz. Patterned bursts of CAPs/CMAPs (vii) recorded from subject NE. The largest of the five bursts in this 1 sec trace have amplitudes of about 0.5 mV. The middle burst in (vii) is detailed in (viii). This 150 msec trace shows that a single burst is a complex of individual CAPs/CMAPs. An EMG (ix) from subject NE during ankle-foot flex one-third through this 2.5 sec trace. Maximum amplitude is about 0.2 mV. (E) Raw human subject recordings highlighting a spike in electrical activity following foot-ankle flexion (black line).

Upon insertion into the skin, we confirmed the DEN device recorded electrical signals from nerves in human subjects without the interference of heart EKG signals (Fig. 6D), which had previously been observed during mouse experiments. There were six major types of neural activity that occurred in human recordings: single action potentials (Fig. 6D, i-ii), spike train activity (Fig. 6D, iii), trains of CAPs/CMAPs (Fig. 6D, iv), individual CAPs/CMAPs (Fig. 6D, v-vi) – which were distinguishable from trains of CAPs/CMAPs, patterned bursts of CAPs/CMAPs (Fig. 6D, vii), and EMG activity following foot-ankle flexion (Fig. 6D, ix). Therefore, we have capability with the DEN to pick up small fiber activity as well as some large fiber activity (EMG in mouse and human). All human recordings were taken at a 40kHz sampling rate, as opposed to the 2kHz sampling rate used in preclinical mouse studies, in order to better visualize individual neural events as seen in Fig. 6D and Suppl. Fig. S3C. Following foot-ankle flexion, a spike in activity across all recording channels was noted (Fig. 6E), which was interpreted as an EMG response. Again, we used our proprietary FOX-DEN software to filter the human DEN recordings of noise and then generated scalograms (Fig. S7A-B), which demonstrated that the DEN was capable of recording nerve electrical activity, including unstimulated action potentials from human subjects.

**Figure 7.**
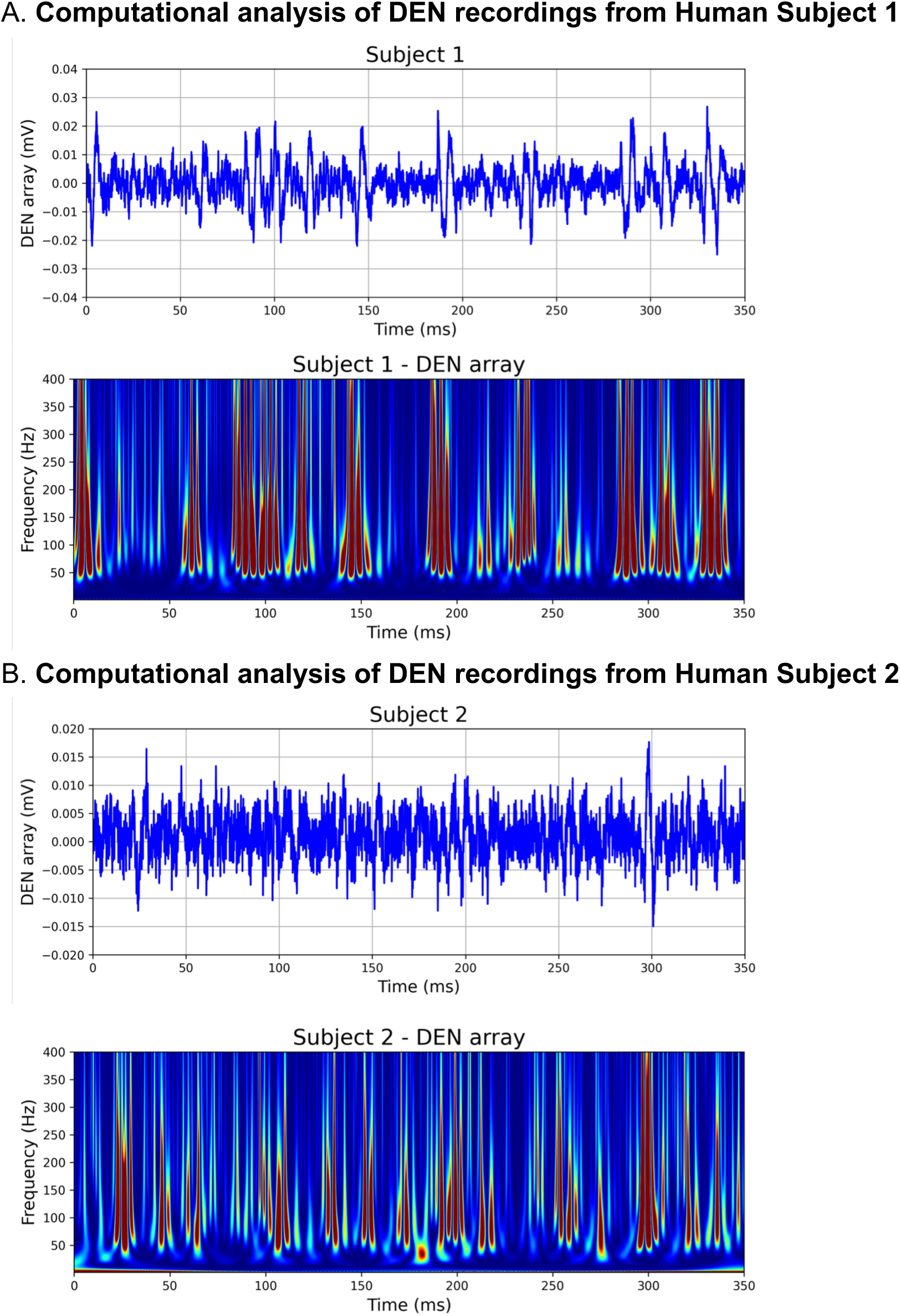
Computationally filtered human DEN recordings. (A) Den recording from Human Subject 1 was computationally filtered using 300Hz 10^th^ order HP Filter (top panel), the resultant scalogram exhibits the signature of a noise-free computed AP train (bottom panel). (B) Computationally filtered DEN recording (top panel) and scalogram (bottom panel) from Human Subject 2.

## Discussion

Following the initial validation studies to evaluate the performance of the DEN device in nerves of invertebrates, we determined that electrical stimulation was not required to record intravital nerve electrical activity from skin and transdermal tissues of control mice. We also confirmed additional settings in the early testing that were used in the diet-induced neuropathy model: the recording site (mouse flank/inguinal skin, with hair removal), the confounding cooling of the animal due to the inhaled anesthetic over time, and the placement of the mouse in a Faraday cage to shield from room electrical noise (which was further filtered out as a reliable 60Hz room noise signal with the BioPac software). This process enabled the progression into a pre-clinical mouse model of DPN, with the goal of determining if the device was able to detect changes to nerve health and activity/function across the progression to DPN, in comparison to other measures such as von Frey and IENFD.

Over the course of this study, we confirmed that the DEN device could successfully record neural activity from both mice and humans in the basal state. A diet-induced pre-diabetic/diabetic mouse model validated that the DEN device could detect pathophysiological changes to nerve activity signatures, associated with a worsening state of diabetes and neuropathy (as confirmed by us through a reduction of paw skin IENFD across the diabetic time-course, and in prior studies by decreased response to noxious stimuli (von Frey)^26^, and reduced NCV at 16wks of diet^25^). In mice, there were three major types of nerve electrical activity seen to occur during the recording process: spike trains, bursts, and single spike events. The activity measurements from the pre-clinical mouse recordings were confirmed to be of neural origin due to the duration of the signals, amplitude, and relative symmetry above and below the y-axis. Specifically, signals of ∼1 msec with a 2:1 signal to noise ratio, symmetry above and below the y-axis, and occurring on only 1 channel at any given time were identified as possible nerve signals. These and other spontaneous field potential electrical events in our recordings likely correspond with various types of neural activity generated within the subdermal tissue and muscle of the mice while under anesthesia, including from the various nerve endings innervating the tissue (sensory, motor, autonomic) at different distances from each of the 4 needle pairs. While these observations were made by the human eye, we also successfully employed unbiased statistical and computational approaches that confirmed these nerve electrical activity signatures were likely of diagnostic value, as they could predict the diet-group (i.e: the extent of DPN) with high accuracy. In fact, in our computational analyses we could confirm the action potential signature, and could tell that some firing was closer to the needles, while others were further away.

By recording with both the DEN device and an EKG electrode, we discovered that the rate of spike trains previously analyzed in mice was synchronous with the rate of EKG over time, and was ruled out as a small fiber nerve signal. Regardless, the ability of the DEN device to record EKG in mice may have beneficial implications when assessing patterns of EKG over a diabetic time course, such as for the diagnosis of cardiac autonomic neuropathy (CAN), and the EKG signal can be computationally filtered out using newer generation FOX-DEN software. One of the advantages of the DEN array is that it captures a much stronger signal in the millivolt range compared with the much weaker signal captured by a surface electrode in the microvolt range (Fig. 4A top panel). This provides a significantly better signal-to-noise ratio for the DEN device compared with a surface electrode device. Interestingly, the rate of spike trains attributed to EKG decreased at 2 weeks of the HFND, a direct correlation to the expected onset of leptin resistance^37^. Leptin has been associated with the moderation of cardiac electrical properties, via inhibition of the β-adrenergic receptor, which can cause leptin to accumulate in the myocardium and result in sustained bradycardia^38^. A similar decrease in spike train/EKG signaling rate occurred in the recordings taken from mice in the 25-week HFND time study. This change may have indicated the presence or development of CAN, a complication associated with diabetes that is often linked to an increased risk of cardiovascular mortality rate^39^. Anesthesia may have also played a role in the depression of heart rate across each recording group over time, but this was controlled for in later assessments and was a convenient added measure in DEN recordings^40^.

One notable adjustment that was made during the transition from recording in mice to recording in humans was the sampling rate at which recordings were obtained. For the mouse recordings, the sampling rate was set at 2kHz, which allowed larger neural and/or muscular events such as CAPs/CMAPs and EMGs to be visualized and evaluated. Recording at 2kHz also managed the file sizes we needed to save and transfer for analyses. However, when recording in humans, the sampling rate was adjusted to 40kHz to better visualize single action potentials alongside larger neural and/or muscular events. Since the sample size in the human data was much smaller than the mouse studies, having a larger file did not pose any problems for analysis at 40kHz. Custom software and cloud data storage solutions are now being developed for future human testing at 40kHz, including for a new clinical trial with the DEN measurements that is now underway.

It is important to note that the DEN recordings presented in this study are not specific to any particular nerve fiber type, nor did we aim to record from a specific subset of nerves. We were intentionally agnostic to which nerve fiber types were being recorded by the DEN, as our aim was to capture electrical activity from any and all nerve fibers potentially affected by polyneuropathy. Electrical signatures from all small nerve fiber types could be captured by the DEN device, and not only allowed us to computationally distinguish a normal versus neuropathic signal, but also showed distinct clustering of electrical signatures as early as 2 wks into a HFND (Fig. 5). Future experiments will focus on identifying critical intervention time points in SFN develop.

Due to the spatial placement of the array, there are multiple potential sources of action potentials, leading to the recording of volume-conducted field potentials, similar to a brain electroencephalogram (EEG)^21^. This type of physiological filtering through tissue can dramatically alter the appearance of an action potential^41^. Volume conduction can change the pattern or shape of an action potential^41^, as reflected in our data; however, our computational filtering of DEN recordings can account for this and present a stereotypical action potential, as shown in Suppl. Fig. S4.

### Limitations

A limitation of our preclinical validation studies was the necessary use of anesthesia to obtain DEN recordings in mice, which resulted in anesthesia-induced suppression of peripheral nerve electrical activity. Although we kept mice at low levels of sedation, obesity affects pharmacokinetics/pharmacodynamics of most anesthetics including isoflurane^42^, and it is unknown to what extent this may have affected nerve electrical activity. It was also clear that the mice DEN recordings had a contaminating, although perhaps useful, EKG signal, but this was not observed in humans and can be filtered out of mouse data using new FOX-DEN software. Finally, mouse skin and subdermal tissues are more densely innervated than human skin and subdermal tissues, as we have observed with our own measurements, and this may contribute to differences in recordings as we expand our human data set.

## Conclusions

In summary, for this study we have validated the functionality of the DEN as a novel diagnostic for SFN/DPN through recording neural activity in the subdermal tissue of basal mice as well as a diet-induced obese/diabetic/neuropathic mouse model. The differences we observed in rates of activity across each time point of HFND support the idea that changes in frequency and/or rate of specific types of nerve activity patterns could indicate pathophysiological changes associated with the onset of early versus late-stage DPN. Our statistical and computational analyses indicated that our recording data were up to 96% accurate in predicting extent of DPN, and that we were measuring changes to nerve electrical activity, such as action potentials in the transdermal recordings. In addition, the DEN was optimized and validated in healthy humans to demonstrate the translational potential of this novel diagnostic approach. These findings support the functionality of the DEN and maintain the possibility that the DEN should be able to provide an earlier and more sensitive diagnosis of DPN and SFN for human patients in the near future.

## Acknowledgements

The authors wish to thank Drs. Christopher Gibbons, Eva Feldman, Sandra Rieger, Brett Morrison, William David Arnold, Miriam Freimer, Bakri Elsheikh, and Ben Harrison for helpful discussions; and Sarrah Marcotte and Caleb Berry for technical assistance in data acquisition and 3D printing. We thank Jonathan Donnelly for early iterations of the statistical design. We particularly want to thank our valued colleague Dr. Jan Pelletier for key insights regarding human skin related to this device development, as well as other helpful discussions and feedback. Dr. Pelletier was a stellar clinician, researcher, colleague, and supporter of this project, and her loss is deeply felt by this research team.

## Author’s Contributions

KLT conceived of the study and oversaw data collection, co-designed the device, and oversaw and contributed to manuscript preparation. RLS led the biomedical engineering team, co-designed the device, and contributed to manuscript preparation. MB led the research team, co-designed the device, analyzed data, and wrote the manuscript. LC collected data and wrote the manuscript. BV, JP, JMT, NS, ER, collected or analyzed data, or contributed to device fabrication and testing. BV also contributed to writing the manuscript. NE oversaw and consulted on the electronics, and LK oversaw and conducted the electrophysiological measurements and analyzed/interpreted data. FK analyzed and modeled the data for statistical learning, and also developed methods to visualize compound action potential trains in DEN data.

## Funding

This project was funded by University of Maine MIRTA and RRF funds, an NSF STTR Phase 1 to Neuright, Inc. (ID: 2014779), Maine Technology Institute (MTI) Seed and Grant funding, Neuright Inc. R&D funding, and an NIH COBRE award from Maine’s Pilot Project Funding Driven program. Julia Towne and Brooke Villinski were supported by NSF REU Site: Sensor Science and Engineering grants 1460700 and 1851998, respectively.

## Ethical Approval and Consent to Participate

All procedures and handling of animals were performed in accordance with the University of Maine’s Institutional Animal Care and Use Committee (IACUC), to comply with the guidelines of the PHS Policy on Humane Care and Use of Laboratory Animals, and Guide for the Care and Use of Laboratory Animals. This study was approved by the University of Maine’s IACUC, under protocol A2020-07-05.

## Availability of Data and Materials

All data can be made available upon request to the corresponding authors.

## Conflicts of Interest

Authors declare no Competing Non-Financial Interests, but the following Competing Financial Interests remain: KLT and MB are co-founders of Neuright, Inc., a University of Maine spin-out and biotechnology R&D start-up company based in Maine and Ohio, which co-funded this study. Neuright, Inc. holds an exclusive license agreement with UMaine to continue R&D and commercialization for this patent-pending theragnostic device platform and associated software, and will conduct future human clinical studies. KTL, MB, and RLS are also named as inventors on a patent filed by UMaine pertaining to the device described in this manuscript: Patent Application: #17/923,293; Status: US publication date 2023-07-27; Application Undergoing Preexam Processing 2024-04-24.

## Abbreviations

CAN: Cardiovascular autonomic neuropathy
CAP: Compound action potential
CMAP: Compound muscle action potential
DEN: Detecting early neuropathy
DPN: Diabetic peripheral neuropathy
EKG: Electrocardiogram
EMG: Electromyography
HFND: High fat neuropathy diet
HPF: High pass filter
IENFD: Intra-epidermal Nerve Fibers
LPF: Low pass filter
PBC: Printed circuit board
PN: Peripheral neuropathy
PAP: Putative action potential
SFN: Small fiber neuropathy
scWAT: Subcutaneous white adipose tissue
SiLN: Subiliac lymph node
WAT: White adipose tissue

## Notes

### Summary of Updates

Additional data (two new figures, including computational analyses) has been added to further validate the DEN device in its ability to pick up small fiber action potentials.

