## Supplemental Figures for "Transdermal Electrophysiological Recordings of Diabetic Small Fiber Peripheral Neuropathy Using a Needle Electrode Array in Mice and Men"

Supplemental Figure S1

**Device validation studies in invertebrates and control/healthy mice**

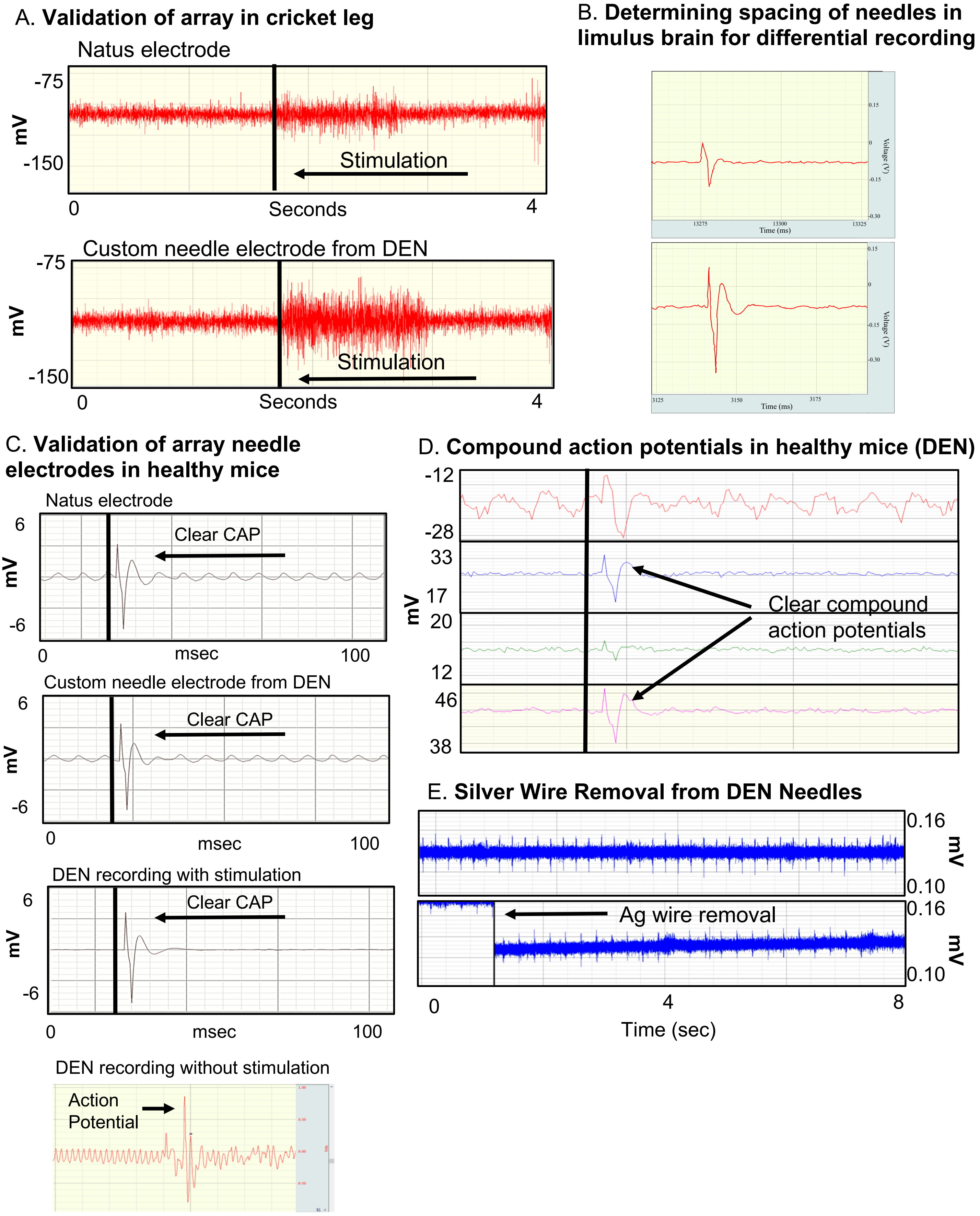

Supplemental Figure S2

A. **Features of spontaneous electrophysiological recordings in mice**

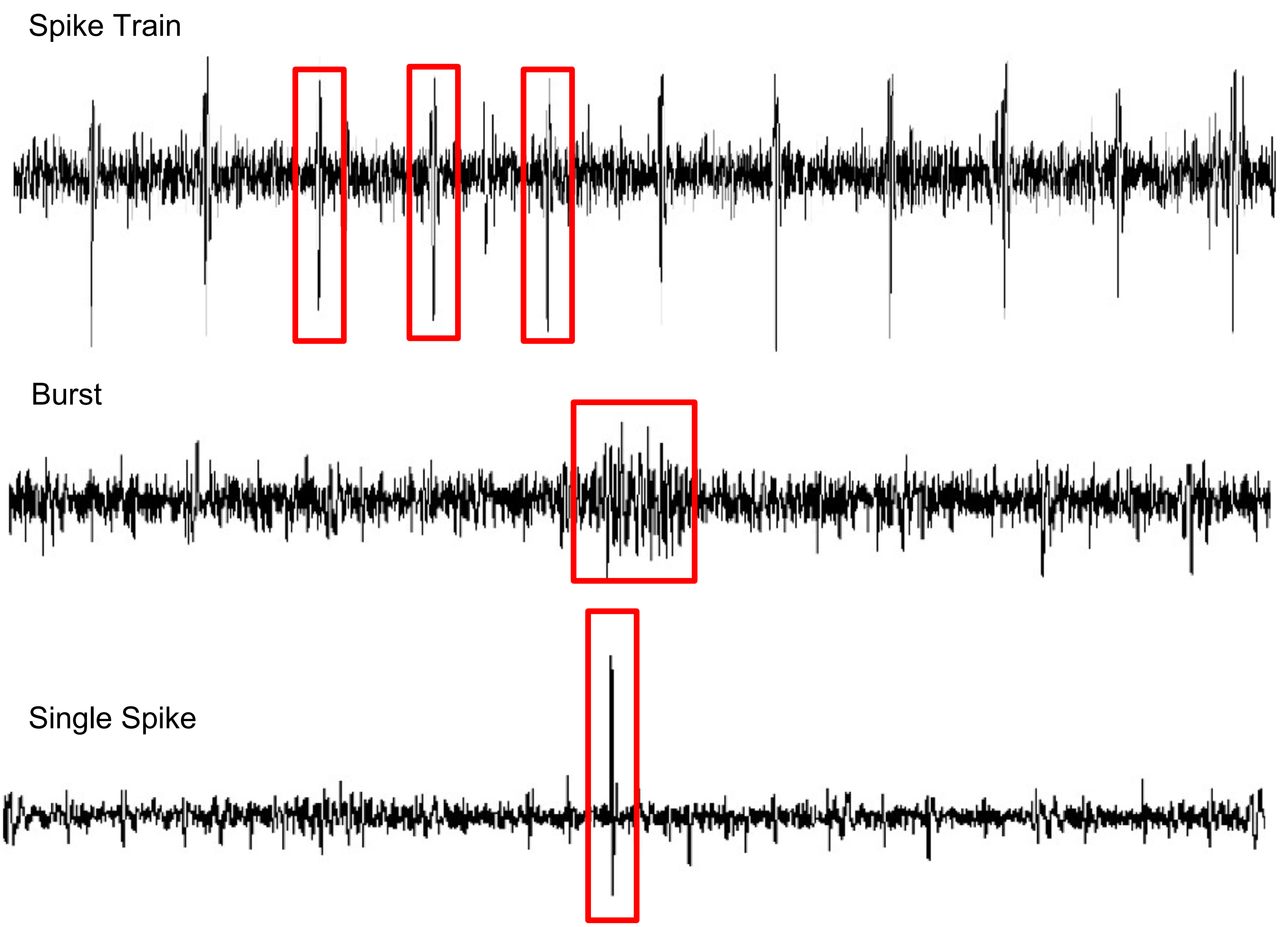


B. **Anatomy of a spike**

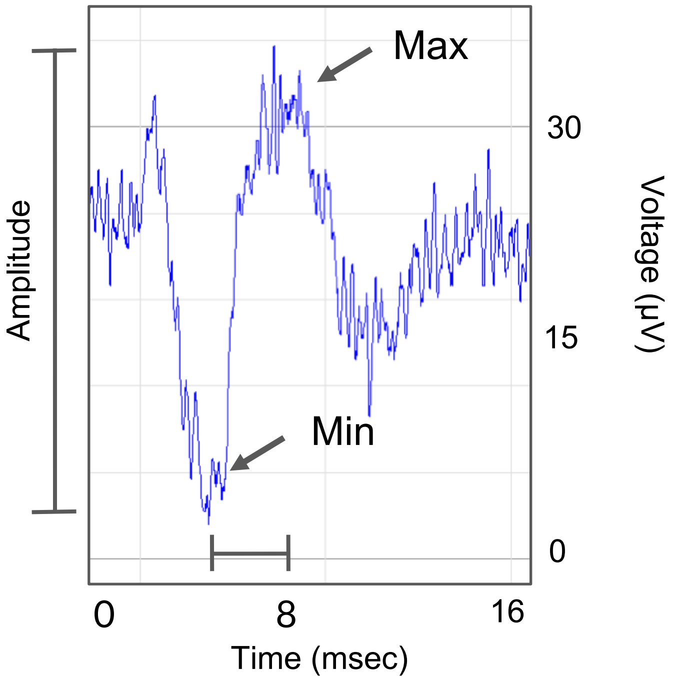

Supplemental Figure S3

A. **Rate of spike occurrence vs. time under anesthesia**

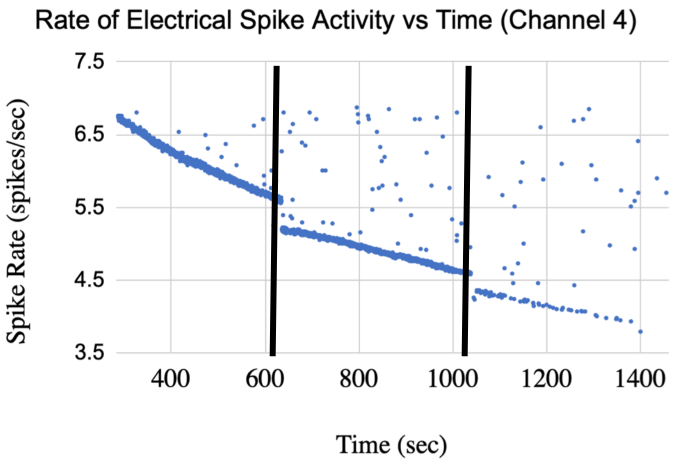


B. **Electrical responses recorded by DEN after warm temperature**

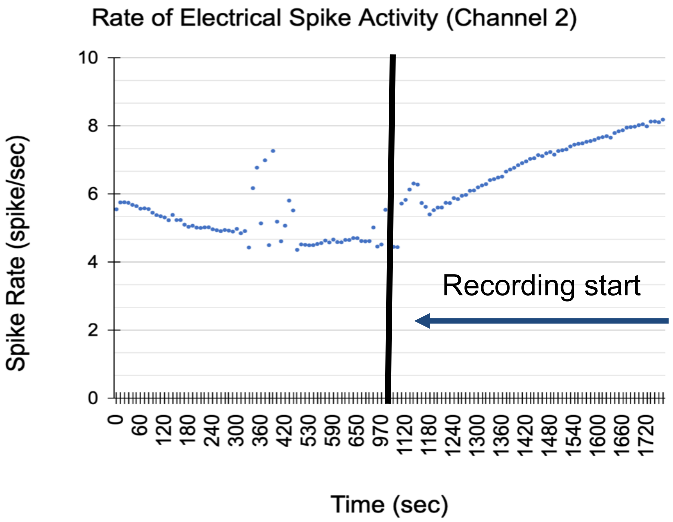


C. **Comparing BIOPAC sampling rate of 2kHz vs. 40kHz with DEN recordings**

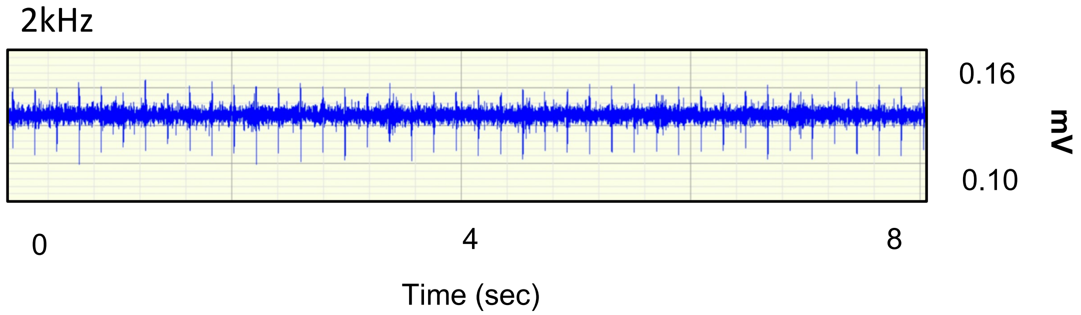


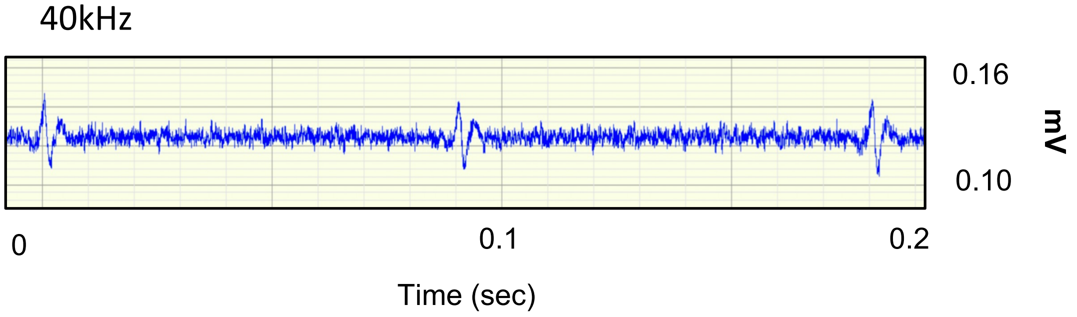

Supplemental Figure S4

A. **Simulated action potential**

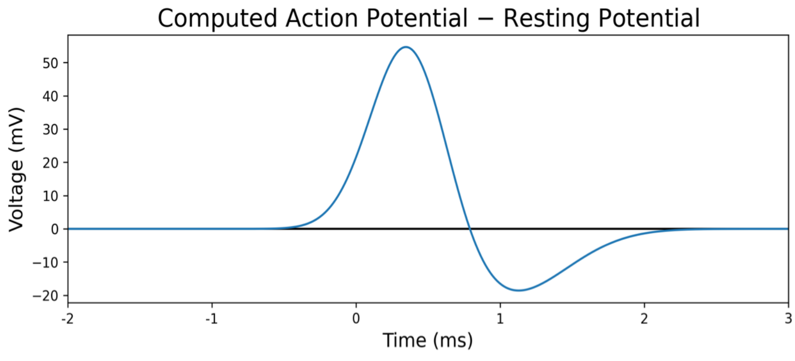


B. **Scalogram contours of simulated action potential train**

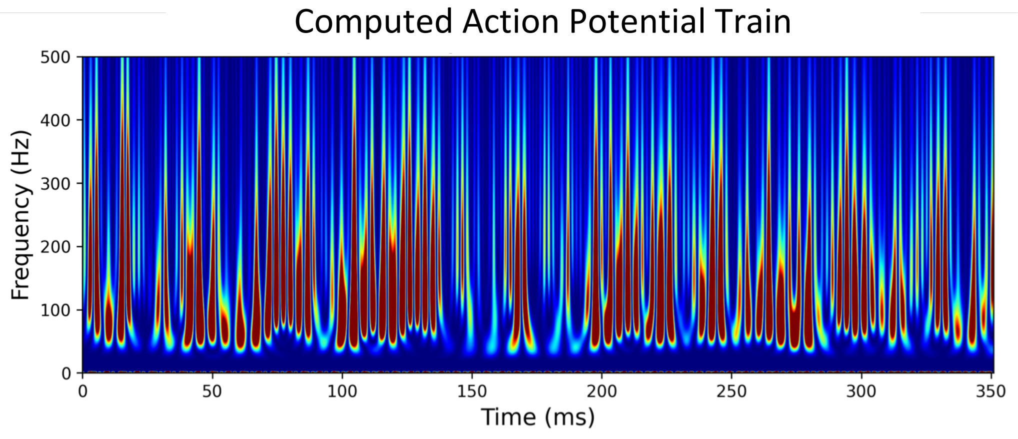


C. **Scalogram contours of Gaussian Noise**

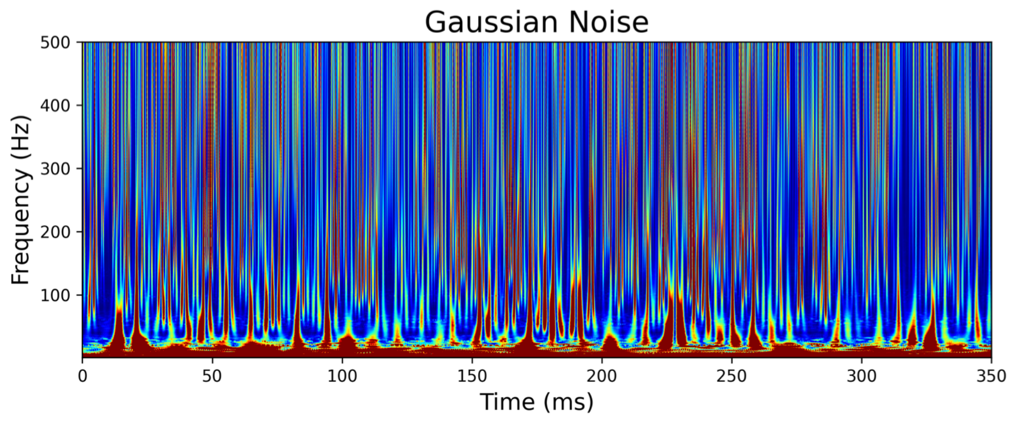

**Supplemental Figure S1. Preliminary validation studies of the DEN device.** (A) Validation of the DEN array vs. Natus electrode in cricket leg, recorded over 4 seconds. Black line denotes onset of mechanical stimulation. (B) Validation of the DEN array vs. Natus electrode vs. whole DEN array in healthy mice. Black line denotes onset of stimulation prior to CAP. (C) Compound action potentials in healthy mice. Black line indicates onset of mechanical stimulation prior to CAP (top 3 panels). Action potential captured by DEN array inserted into flank of healthy mouse (bottom panel). (D) Compound action potentials captured by the DEN in a healthy mouse following stimulation. (E) Removal of the silver wire inside the needle electrodes showed no significant difference in the recording sensitivity of DEN, as shown by the continuation of a spike train after wire removal (black line).

**Supplemental Figure S2: Features of electrophysiological recordings obtained by the DEN in mice.** (A) The 3 major types of activity recorded from mice: spike train activity (duration of 2msec shown), burst activity (duration of 1msec shown), and single spike (duration of 1msec shown). Baseline noise for traces were approximately 20mV. (B) Anatomy of a possible action potential (PAP) consisting of a single spike.

**Supplemental Figure S3. Physiological validation of recordings and sample rate.** (A) Rate of spike activity occurrence over time recorded from a high fat diet animal. Black lines denote the repositioning of DEN. (B) Hot water bottle experiment with graph showing rate of spike train activity over time, with each point denoting an instantaneous rate of activity. Black line indicates start of recording after heating the mouse with a hot water bottle for approximately 5 minutes. (C) 2kHz vs 40kHz mouse recording showing difference in recording sensitivity. Both recordings taken from a HFND animal.

**Supplemental Figure S4. DEN data is mostly composed of AP trains with low noise levels.** (A) Theoretical simulation of an action potential. (B) The scalogram of a computed action potential train. (C) Scalogram of signatures of random noise for comparison.
